# Genetic liability for internalizing versus externalizing behavior manifests in the developing and adult hippocampus: Insight from a meta-analysis of transcriptional profiling studies in a selectively-bred rat model

**DOI:** 10.1101/774034

**Authors:** Isabelle A. Birt, Megan H. Hagenauer, Sarah M. Clinton, Cigdem Aydin, Peter Blandino, John D. H. Stead, Kathryn L. Hilde, Fan Meng, Robert C. Thompson, Huzefa Khalil, Alex Stefanov, Pamela Maras, Zhifeng Zhou, Elaine K. Hebda-Bauer, David Goldman, Stanley J. Watson, Huda Akil

## Abstract

**Background:** For over 16 years, we have selectively bred rats to show either high or low levels of exploratory activity within a novel environment. These “bred High Responder” (bHR) and “bred Low Responder” (bLR) rats serve as a model for temperamental extremes, exhibiting large differences in many internalizing and externalizing behaviors relevant to mood and substance abuse disorders.

**Methods:** Our study elucidated persistent differences in gene expression related to bHR/bLR phenotype across development and adulthood in the hippocampus, a region critical for emotional regulation. We meta-analyzed eight transcriptional profiling datasets (microarray, RNA-Seq) spanning 43 generations of selective breeding (adult: *n*=46, P7: *n*=22, P14: *n*=49, P21: *n*=21; all male). We cross-referenced these results with exome sequencing performed on our colony to pinpoint candidates likely to mediate the effect of selective breeding on behavioral phenotype.

**Results:** Genetic and transcriptional profiling results converged to implicate two genes with previous associations with metabolism and mood: Thyrotropin releasing hormone receptor and Uncoupling protein 2. Our results also highlighted bHR/bLR functional differences in the hippocampus, including a network essential for neurodevelopmental programming, proliferation, and differentiation, containing hub genes Bone morphogenetic protein 4 and Marker of proliferation ki-67. Finally, we observed differential expression related to microglial activation, which is important for synaptic pruning, including two genes within implicated chromosomal regions: Complement C1q A chain and Milk fat globule-EGF factor 8.

**Conclusion:** These candidate genes and functional pathways have the capability to direct bHR/bLR rats along divergent developmental trajectories and promote a widely different reactivity to the environment.

## Introduction

The strong pattern of comorbidity amongst psychiatric disorders is believed to be generated by a spectrum of latent liability (1), arising from a complex interplay of genetic risk and environmental factors, such as stress and childhood adversity (1–3). At one end of this spectrum are internalizing disorders, which are associated with neuroticism, anxiety, and depression. At the other end of the spectrum are externalizing disorders, which are associated with risk-taking and novelty-seeking, as seen in mania, substance abuse, and impulse-control disorders (1).

We model the genetic contributions underlying both extremes of this spectrum by selectively breeding rats that react differently to a novel environment. “Bred High Responder” (bHR) rats are highly exploratory with a disinhibited, novelty-seeking temperament, including hyperactivity, aggression, and drug-seeking. “Bred Low Responder” (bLR) rats are highly-inhibited, exhibiting reduced locomotor activity and anxious and depressive-like behavior (4–13). These behavioral propensities are robust and stable, beginning early in development (14, 15) similar to temperament in humans (16).

This highly-differentiated phenotype makes bHR/bLR rats ideal for observing the developmental programming and adult manifestation of neurological factors underlying internalizing and externalizing tendencies (8,11,17). This study focused on the hippocampus, a region important for emotional regulation, behavioral inhibition (18–20), and reactivity to the environment (18), including stress-related glucocorticoid release (21, 22). In the bHR/bLR model, we previously observed differences in hippocampal glucocorticoid receptor and growth factor expression, histone methylation, cell proliferation, survival, and overall volume (5,9,14,23,24).

Our current study characterized hippocampal gene expression in bHR/bLR rats across development and adulthood using a meta-analysis of eight transcriptional profiling datasets spanning 43 generations of selective breeding. Concurrently, we discovered chromosomal regions containing bHR/bLR segregating variants that are likely to contribute to exploratory locomotor phenotype (25). By comparing across these studies, we identified differentially-expressed (DE) genes situated within implicated chromosomal regions and hippocampal functional pathways essential for mood and development (Fig 1). These genes are promising candidates for mediating the influence of selective breeding on behavioral phenotype.

**Figure 1.**
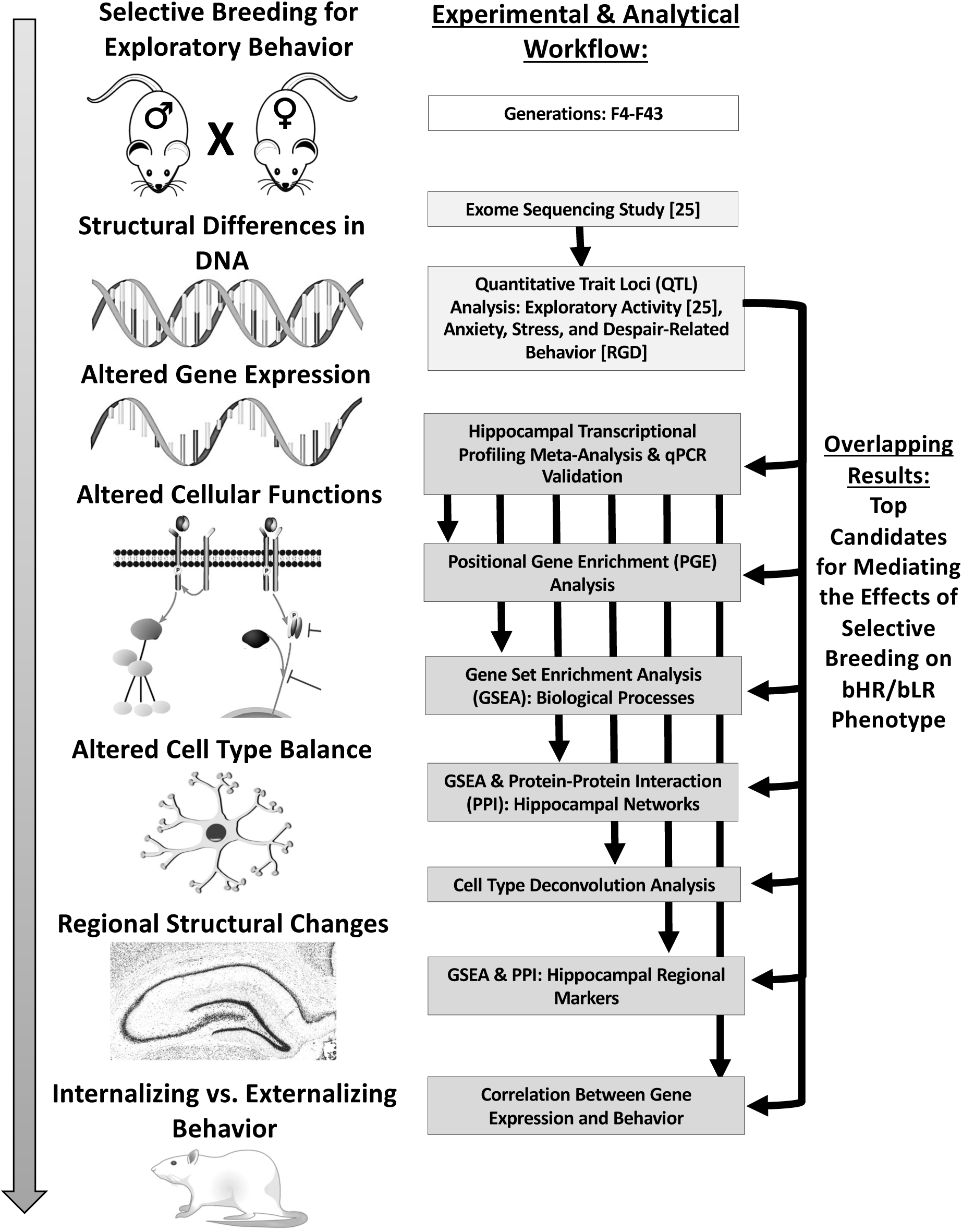
An overview of the experimental and analytical workflow used to identify top candidate genes for mediating the effects of selective breeding on bHR/bLR phenotype. ***Left:*** Many generations of selective breeding based on exploratory locomotor behavior drove segregation of genetic variants that contribute to internalizing and externalizing behavior within our bHR and bLR rats. The effect of these variants on behavior is mediated by alterations in gene expression and cellular function, which produce local changes in cell type balance and structure within brain regions responsible for affective behavior, such as the hippocampus. ***Right:*** Our concurrent genetic study used exome sequencing to identify genetic variants that segregated bHR/bLR rats in our colony, and then used a sampling of those variants to locate regions of the genome (quantitative trait loci – QTLs) that might contribute to exploratory locomotor activity (25). Our current study used meta-analyses of hippocampal transcriptional profiling studies to identify bHR/bLR differentially-expressed (DE) genes, pathways, cell types, and networks in development and adulthood. In our results, we emphasize DE genes that were *1)* consistently DE across multiple developmental stages, *2)* central to DE pathways, cell types, and networks, *3)* located near genetic variants that segregated bHR/bLR rats in our colony and/or within QTLs for exploratory locomotion. These genes are the top candidates for mediating the effects of selective breeding on bHR/bLR phenotype.

## Methods

Full methods for the individual experiments and analyses are provided in the supplement. The associated datasets have been released on GEO (Table 1) or FigShare (https://doi.org/10.1101/774034). Analysis code (R Studio v.1.0.153, R v.3.2.2) is available at https://github.com/isabellie4/PhenotypeProject and https://github.com/hagenaue/bHRbLR_MetaAnalysisProject.

**Table 1.** An overview of the eight transcriptional profiling studies included in our current meta-analyses of differential gene expression in the bHR and bLR hippocampus at four developmental time points: P7, P14, P21, and adulthood. Citations: (28, 125)

|  | MBNI_AffymetrixRae230_F4 | MBNI_AffymetrixRgU34A_F6 | MBNI_AffymetrixRae230_F15 | MBNI_IlluminaRatRef12v1_F15 | MBNI_RNASeq_F29 | Alabama_Nimblegen_F34 | MBNI_RNASeq_F37 | MBNI_RNASeq_F43 |
| --- | --- | --- | --- | --- | --- | --- | --- | --- |
| **Generation** | 4 | 6 | 15 | 15 | 29 | 34 | 37 | 43 |
| **Laboratory** | MBNI | MBNI | MBNI | MBNI | MBNI | Alabama | MBNI | MBNI |
| **Lead Scientist** | Dr. John Stead | Dr. Sarah Clinton | Dr. Sarah Clinton | Dr. Sarah Clinton | Dr. Sarah Clinton | Dr. Sarah Clinton | Dr. Peter Blandino | Dr. Cigdem Aydin |
| **Age** | Adult | P7, P14, P21 | P14 | P14 | P14, Adult | P7, P14, P21, Adult | Adult | Adult |
| **n per group** | 6 | 6 | 6 | 6 | 2 | 5 | 6 | 5 |
| **Platform** | Affymetrix Rat Expression Set 230 A | Affymetrix Rat Genome (RG) U34A GeneChips | Affymetrix Rat Expression Set 230 A | Illumina RatRef-12v1 Beadchip | RNA-Seq | NimbleGen Rat Gene Expression 12x135 | RNA-Seq | RNA-Seq |
| **Tissue Extraction** | Whole Hippocampus | Whole Hippocampus | Whole Hippocampus | Whole Hippocampus | Whole Hippocampus | Dorsal Hippocampus Tissue Punch | Whole Hippocampus | Whole Hippocampus |
| **Exposure** | Basal | Basal | Basal | Basal | Basal | Basal | Basal | Vehicle Injections |
| **Behavioral Testing** | Basal | Basal | Basal | Basal | Basal | Basal | Locomotor, Anxiety | Social Interaction After Mild Stress |
| **GEO Data Release** | GSE140596 | GSE29552 | GSE140595 | GSE140594 | GSE140597 | GSE88874 | GSE140598 | GSE140287 |
| **Citation** |  | Clinton *et al.* 2011 |  |  |  | Cohen *et al.* 2015 |  |  |

### The bHR/bLR Rat Colony

All experiments were approved by the local University Committee on the Use and Care of Animals, in accordance with the National Institutes of Health Guide for the Care and Use of Laboratory Animals.

#### Selective Breeding

We began selectively-breeding bHR/bLR rats in the Molecular Behavioral Neuroscience Institute (MBNI) at the University of Michigan in 2003 (protocol: (11)). Later, a second colony was begun at the University of Alabama-Birmingham using generation F30 bHR/bLR rats from MBNI. Our meta-analyses use datasets derived from male bHR/bLR rats spanning generations F4-F43. We refer to these datasets according to their respective institution, transcriptional profiling platform, and generation (Table 1). *MBNI_RNASeq_F37* also included a bHRxbLR cross (“Intermediate Responder” (bIR) rats)).

#### Behavioral Testing

For each generation, locomotor response to a novel environment was assessed between P50–75 (11). For *MBNI_RNASeq_F37*, we measured anxiety-like behavior in adulthood (bHR/bLR: P160-P167; bIR: P65-75) using the percent time spent in the open arms of an Elevated Plus Maze (EPM; 5 min test, protocol: (26)). For *MBNI_RNASeq_F43*, we measured social interaction in adulthood (P92, protocol: (4)) after 15 min exposure to the anxiogenic open arms of the EPM (protocol: (27)).

### Hippocampal Gene Expression Analyses

#### Broad Overview of the Datasets

Our meta-analyses included eight datasets from bHR/bLR rats aged P7, P14, P21, and adult (Table 1). The rats were housed in standard conditions with minimal intervention besides behavioral testing or saline injections, and sacrificed by rapid decapitation without anesthesia. The whole hippocampus was dissected, except in *Alabama_Nimblegen_F34*, where dorsal hippocampal tissue punches were performed on sliced frozen tissue (28). The extracted RNA was profiled using microarray (earlier generations: F4, F6, F15, F34) or RNA-Seq (later generations: F29, F37, F43).

#### Broad Overview of the Data Preprocessing

The pre-processing steps for each study varied according to platform, but included common steps, including re-annotation, normalization to reduce technical variation, and quality control. Microarray data were typically summarized into log2-transformed expression sets using Robust Multi-Array Average (RMA: (29)). Gene-level RNA-Seq read count summaries were converted to log(2) fragments per million (FPM). When applicable, transcript data were averaged by gene symbol to obtain a single expression value per sample per gene.

#### Meta-Analyses

Within each dataset, we calculated the effect size (Cohen’s d and variance of d) of bHR/bLR phenotype on the expression data for each gene within each age group. This output was aligned across datasets using official gene symbol. The meta-analysis for each age group was performed using *rma.mv()* (package *metafor* (30)) and corrected for false discovery rate (FDR; Benjamini-Hochberg method in *multtest*: (31)). Including generation as a co-variate provided little additional insight (*generation*: all genes FDR>0.3).

#### Overlap with Previously-Identified Genetic Variants

Our exome sequencing study identified bHR/bLR segregating variants (single-nucleotide polymorphisms, or SNPs), and used a sampling of those variants to pinpoint quantitative trait loci (QTLs) for exploratory locomotion using an bHRxbLR F2 intercross (25). In our current study, we identified all DE genes nearby (+/− 1 MB) the segregating variants and QTL peaks (LOD>3) and determined overlap with additional QTLs relevant to bHR/bLR behavioral phenotype from Rat Genome Database ((32), accessed 08/08/2019, keywords: “Anxiety”, “Stress”, and “Despair”).

#### Positional Gene Enrichment Analysis

To explore which genes might be either co-regulated or in linkage disequilibrium with a causal genetic variant, we evaluated the clustering of top bHR/bLR DE genes within chromosomal regions using Positional Gene Enrichment analysis (*PGE*, http://silico.biotoul.fr/pge/, (33)) and the top results from the P14 and adult meta-analyses (nominal p<0.01, removing duplicated EntrezIDs).

#### Gene Set Enrichment Analysis (GSEA) & Protein-Protein Interaction (PPI) Networks

To elucidate overall functional trends within the largest sets of results (P14 and Adult) we used GSEA ((34, 35); package *fgsea* (36)) and gene set matrix files (.gmt) containing standard gene ontology for rats (go2msig.org; (37)), or customized hippocampal-specific gene sets **(Table S1)** including previously-identified co-expression modules (38, 39) and expression specific to hippocampal neuronal subtypes or subregions (*Hipposeq:* (40)). We further explored top DE genes (adult meta-analysis FDR<0.10) and implicated hippocampal gene sets using predicted PPI networks (string-db.org (41)).

#### Cell Type Data Deconvolution

To interrogate the relative cell type composition of our samples, we used the *BrainInABlender* method (42). Data for genes previously identified as having cell type specific expression was extracted, normalized, and averaged to produce a “cell type index”. For this analysis, we excluded the small MBNI_RNASeq_F29 dataset (n=2/group). We then performed a meta-analysis of the effects of bHR/bLR phenotype on these cell type indices using aforementioned methods.

#### qPCR Validation

Hippocampal tissue from 6 bHR and 6 bLR males was collected at ages P14 (generation F55) and P90 (generation F51; **Fig S1**). Following cDNA synthesis, Bone morphogenetic protein 4 (Bmp4) was quantified using qPCR and custom-designed primers, using the Livak method (43) and glyceraldehyde-3-phosphate dehydrogenase (Gapdh) as reference. Group differences in -ΔC_q_ at each age were assessed using Welch’s two-sample t-test (44).

## Results

### Selective Breeding Amplifies the Propensity for Internalizing vs. Externalizing Behavior

The divergence of bHR/bLR exploratory activity in response to selective breeding happened rapidly (Fig 2A), implying oligogenic inheritance (25). This divergence was accompanied by a general amplification of internalizing and externalizing tendencies (11,14,45,46). For example, in the behavioral data accompanying our transcriptional profiling datasets, bLRs showed more anxiety-like behavior than bHRs (Fig 2B), and spent less time interacting socially following a stressful challenge (Fig 2C). Therefore, we expected that examining gene expression across bHR/bLR generations would reveal a convergence of effects within implicated chromosomal regions and pathways essential to affective behavior and reactivity to the environment.

**Figure 2.**
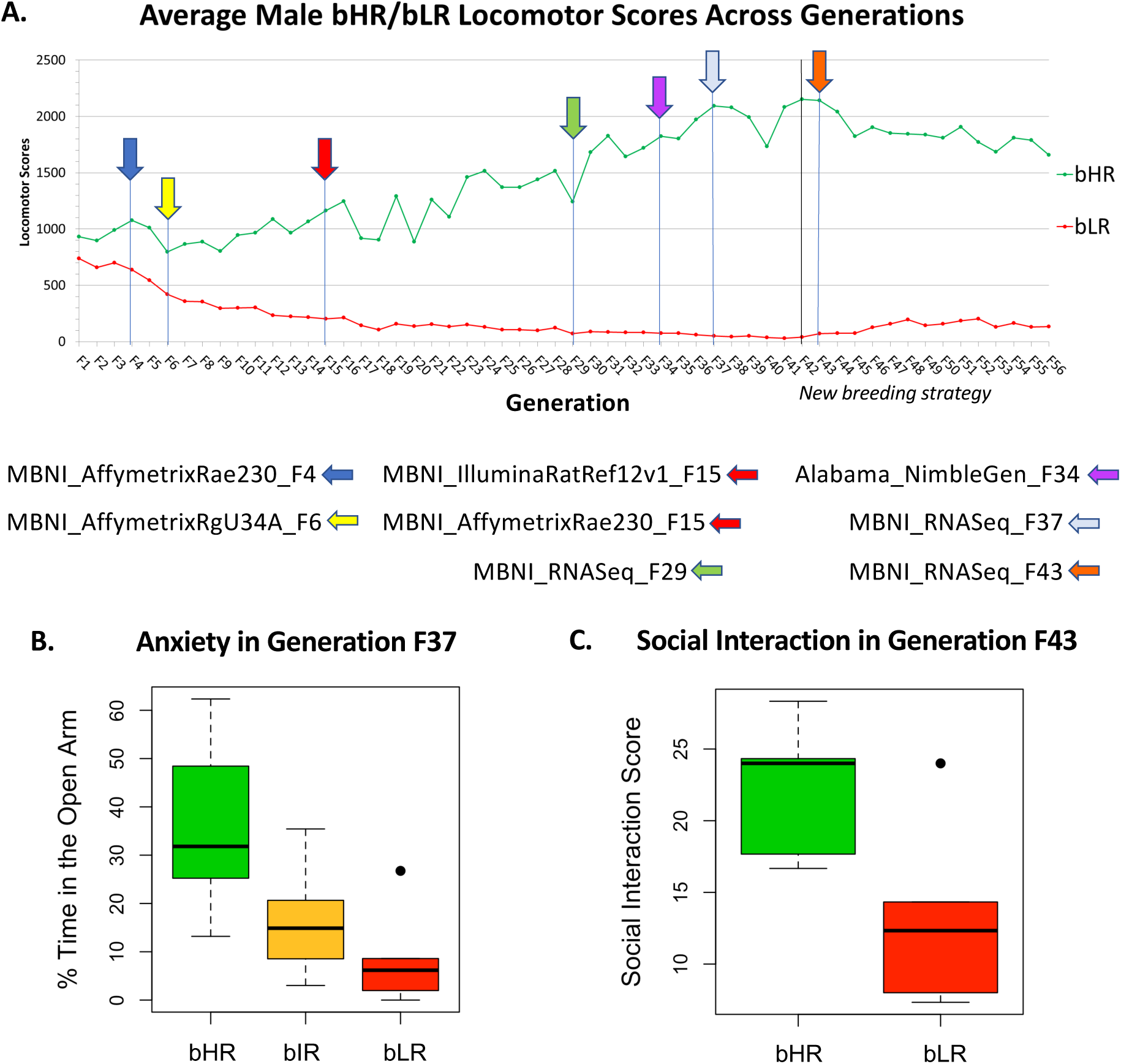
Selectively-bred high responder (bHR) and low responder (bLR) rats model an extreme propensity for internalizing vs. externalizing behavior. **A)** Over the course of 56 generations of selective breeding (F1-F56), the bHR rats (green) have developed increasingly elevated exploratory activity in a novel field (*y-axis:* average total locomotor score), whereas the bLR rats (red) have become less exploratory. These trends plateaued after F42, when our breeding strategy changed to deaccelerate divergence. Arrows indicate the generations during which hippocampal transcriptomic profiling datasets were collected, along with a name indicating the respective laboratory, platform, and generation for each dataset. **B)** bLR rats have been highly anxious since the initiation of our breeding colony. The example above is from the behavioral data accompanying the MBNI_RNASeq_F37 transcriptomic dataset showing bLRs spending a smaller percentage of time in the anxiogenic open arms of the elevated plus maze than bHR rats (effect of phenotype: *F*(2,15)=6.72, *p*=8.25E-03, boxes=first quartile, median, and third quartile, whiskers = range). **C)** bLR rats are more reactive to stressors. This example is from the behavioral data accompanying the MBNI_RNASeq_F43 transcriptomic dataset showing bLR rats spending a smaller percentage of time interacting socially following exposure to a single mild stressor (*F*(1,8)=5.86, *p*=0.0418).

### Selective Breeding for Exploratory Locomotion Alters Hippocampal Gene Expression

Between generations F4-43, we conducted eight exploratory studies transcriptionally profiling the hippocampus of bHR/bLR rats at four ages (P7, P14, P21, and adult). These small studies individually produced few reliable results (**Figs S2-S3**). Nevertheless, a formal meta-analysis revealed multiple genes with consistent DE across generations (Fig 3, **Table S2**). These results can be explored interactively at https://mbni.org/dashboard/huzefak/hrlrMetaAnalysis/.

**Figure 3.**
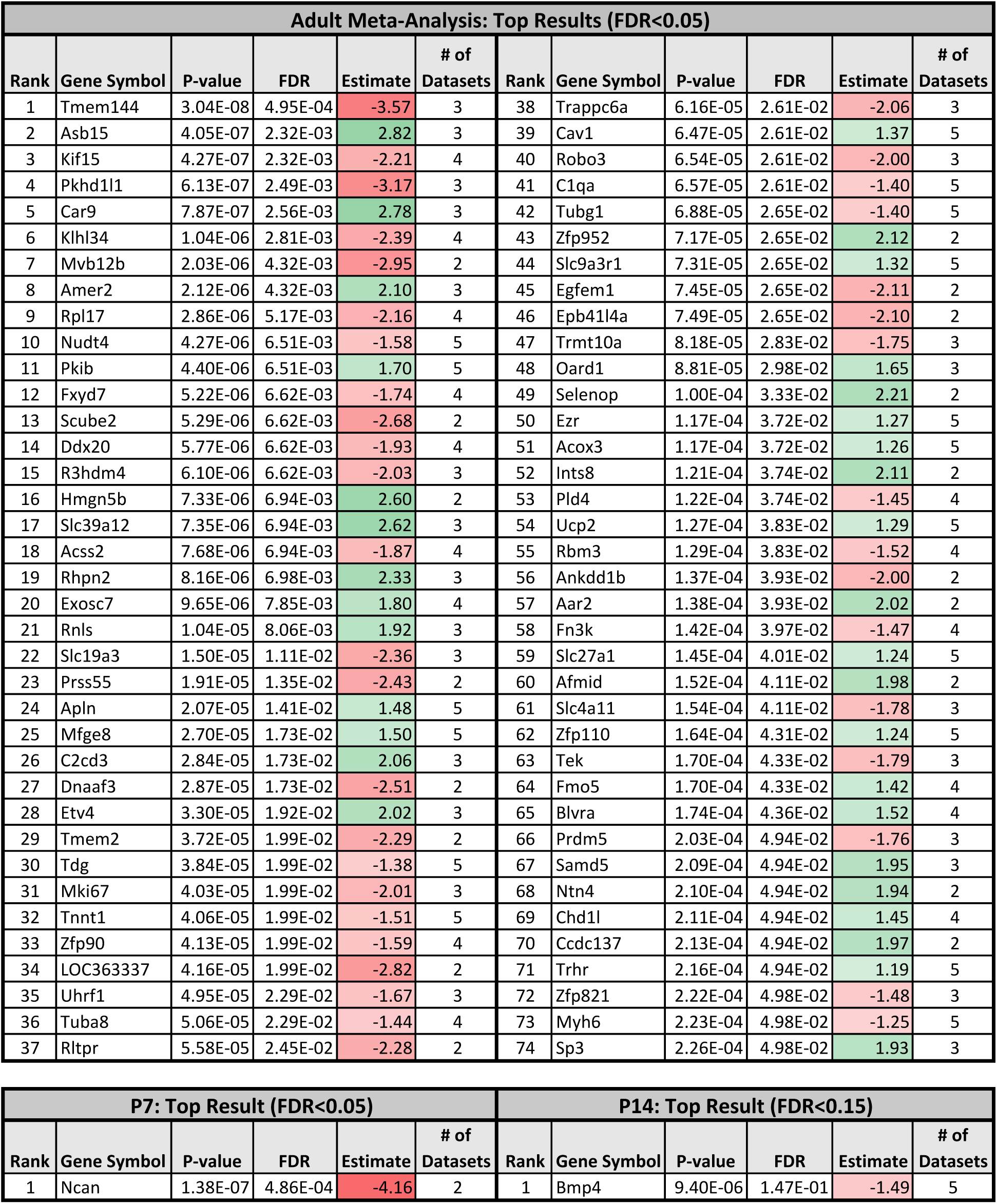
The top DE genes within the bHR/bLR hippocampal gene expression meta-analyses. P-value=nominal p-value, FDR=false detection rate, Estimate=estimated effect size (i.e., the difference in expression between bHR and bLR rats in units of standard deviation, green/positive=higher expression in bHRs, red/negative= higher expression in bLRs), # of datasets=number of datasets included in the meta-analysis for that gene.

#### Adulthood

The effect of bHR/bLR phenotype on gene expression was significant for 74 genes (FDR<0.05, out of 16,269; Fig 3). In general, the estimated effect sizes (*β*’s) tended to be more extreme for genes exclusively represented in RNA-Seq data from later generations (**Fig S4**), but due to smaller sample size, these effects were not overrepresented in the top results. To rule out bias due to platform, we ran a meta-analysis using only recent RNA-Seq data (F37/F43) and confirmed that similar DE genes and pathways were identified (**Fig S5**). In the following discussion we emphasize candidate genes for which there is evidence that expression diverged during the earliest generations.

#### Development

The developmental meta-analyses were more dependent on data from earlier generations and produced less robust results. However, the top genes had consistent effects across age groups. Within the P7 meta-analysis, only one gene (out of 3,257), Neurocan (Ncan), was differentially expressed, with higher expression in bLRs than bHRs since an early generation (Fig 4B-C), in a manner that nominally persisted at P14 (Fig 4D). Within the P14 and P21 meta-analysis, none of the effects survived multiple comparison correction (for 15,682 and 3,257 genes, respectively). However, the top gene at P14, Bmp4, was consistently expressed at higher levels in bLRs than bHRs within both the P14 (Fig 5A) and adult datasets since the F4 generation (Fig 5B). We confirmed DE at these ages using qPCR (Fig 5C, **Fig S1**).

**Figure 4.**
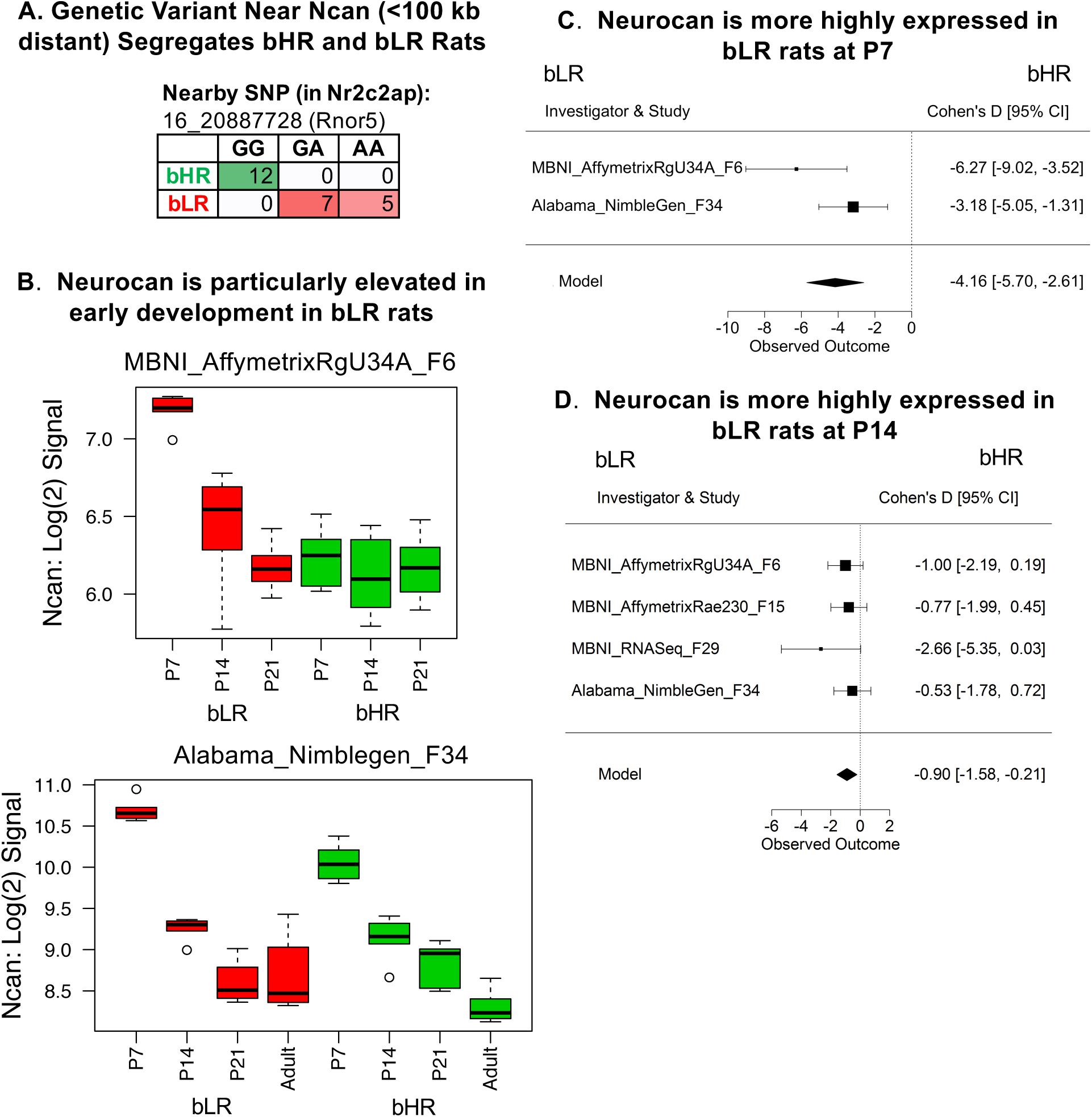
The extracellular matrix constituent Neurocan (Ncan) has elevated expression in bLR rats at age P7. **A)** A genetic variant on Chr 16 nearby Ncan (<100 kb) segregated bHR and bLR rats (Fisher’s exact test: *p*=1.63E-07). **B)** Boxplots illustrating the effect of age and bHR/bLR phenotype on Ncan expression (log(2) signal) in two microarray studies (boxes=median and interquartile range, whiskers=range, red=bLR, green=bHR). The effect of phenotype is most obvious at an age when Ncan is elevated in development (P7). **C-D)** Forest plots showing that Ncan was more expressed in bLRs than bHRs (boxes=Cohen’s D from each study +/-95% confidence intervals, “Model”=estimated effect size +/-95% confidence intervals provided by the meta-analysis) **C)** in the P7 meta-analysis (*β*=-4.16, *p=*1.38E-07, *FDR=*4.86E-04), **D)** and nominally in the P14 meta-analysis (*β*=-0.90, *p=*1.01E-02, *FDR=*7.24E-01).

**Figure 5.**
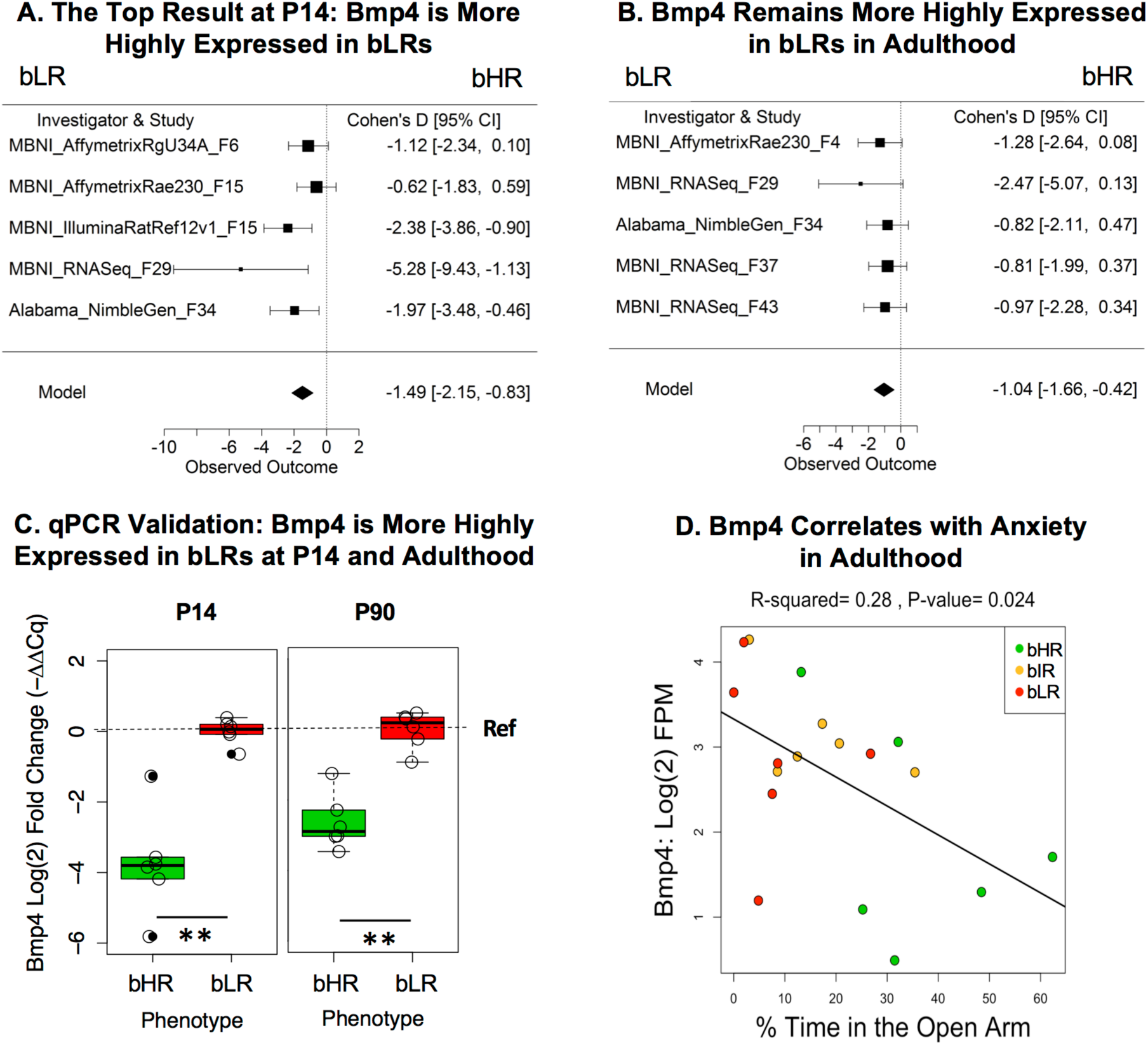
A regulator of proliferation and differentiation, Bone morphogenetic protein 4 (Bmp4), is more highly expressed in bLR rats than bHR rats at P14 and adulthood. **A-B)** Two forest plots showing that Bmp4 was consistently elevated in bLR rats (boxes=Cohen’s D from each study +/-95% confidence intervals, “Model”=estimated effect size +/-95% confidence intervals provided by the meta-analysis) at **A)** P14 (*β*=-1.49, *p*=9.40E-06, *FDR*=1.47E-01) and **B)** adulthood (adult: *β*=-1.04, *p*=1.01E-03, *FDR*=9.38E-02). This direction of effect mirrors findings in the literature that show that blocking the expression of Bmp4 in mice reduces anxiety and depressive-like behavior (98). **C)** Using qPCR, we confirmed that bLRs showed greater *Bmp4* expression than bHRs at P14 and adulthood (P90) using hippocampal tissue from later generations (F51, F55). Log(2) fold change in *Bmp4* expression was calculated using the Livak method (-ΔΔCq, (43)), using *Gapdh* as the reference housekeeping gene and bLRs as the reference group (therefore the bLR mean is set to 0 in all panels). \*\**p*<0.005 (Data release: doi: 10.6084/m9.figshare.10321658; *P14*: *Log(2)FC*=-3.74, *T*(5.60)=-6.10, *p*=0.00115; *P90*: *T*(8.74)=-6.87, *p*=8.44E-05). **D)** Within the behavioral data accompanying the MBNI_RNASeq_F37 dataset, we found that Bmp4 showed a negative relationship with percent time in the open arms (*β*=-0.034, *R^2^*= 0.28, *p*=0.024) and a positive relationship with the number of fecal boli produced on the EPM (*β*=0.32, *R^2^*= 0.29, *p*=0.020, *data not shown*).

### Many bHR/bLR Differentially Expressed Genes Are Located Within Implicated Chromosomal Regions

Our exome sequencing study identified bHR/bLR segregating genetic variants, and then used a sampling of those variants to identify chromosomal regions that are likely to contribute to exploratory locomotor phenotype (QTL peaks). These implicated chromosomal regions overlapped extensively with QTLs relevant to internalizing/externalizing behavior (32), including anxiety (47–52), stress response (53–55), and behavioral despair (56). In our current study, 68% of the genes DE in adulthood (FDR<0.05) were within +/-1 MB of a bHR/bLR segregating variant, and 21% were within +/-100 kB of a highly segregating variant (Bonferonni-corrected *α*=5.00E-05), a 4.7x enrichment compared to non-DE genes (Fig 6A, Fischer’s exact test: p=3.50E-06). DE genes were also 4.5x more likely to be located within a QTL for exploratory locomotion (LOD>4; Fig 6B). This overlap included two genes previously associated with internalizing and externalizing behaviors (Thyrotropin Releasing Hormone Receptor (Trhr), Uncoupling Protein 2 (Ucp2), Fig 6C, (57–62)). These results fit our expectation that the influence of bHR/bLR segregating genetic variants on exploratory locomotion is at least partially mediated by effects on gene expression within the hippocampus.

**Figure 6.**
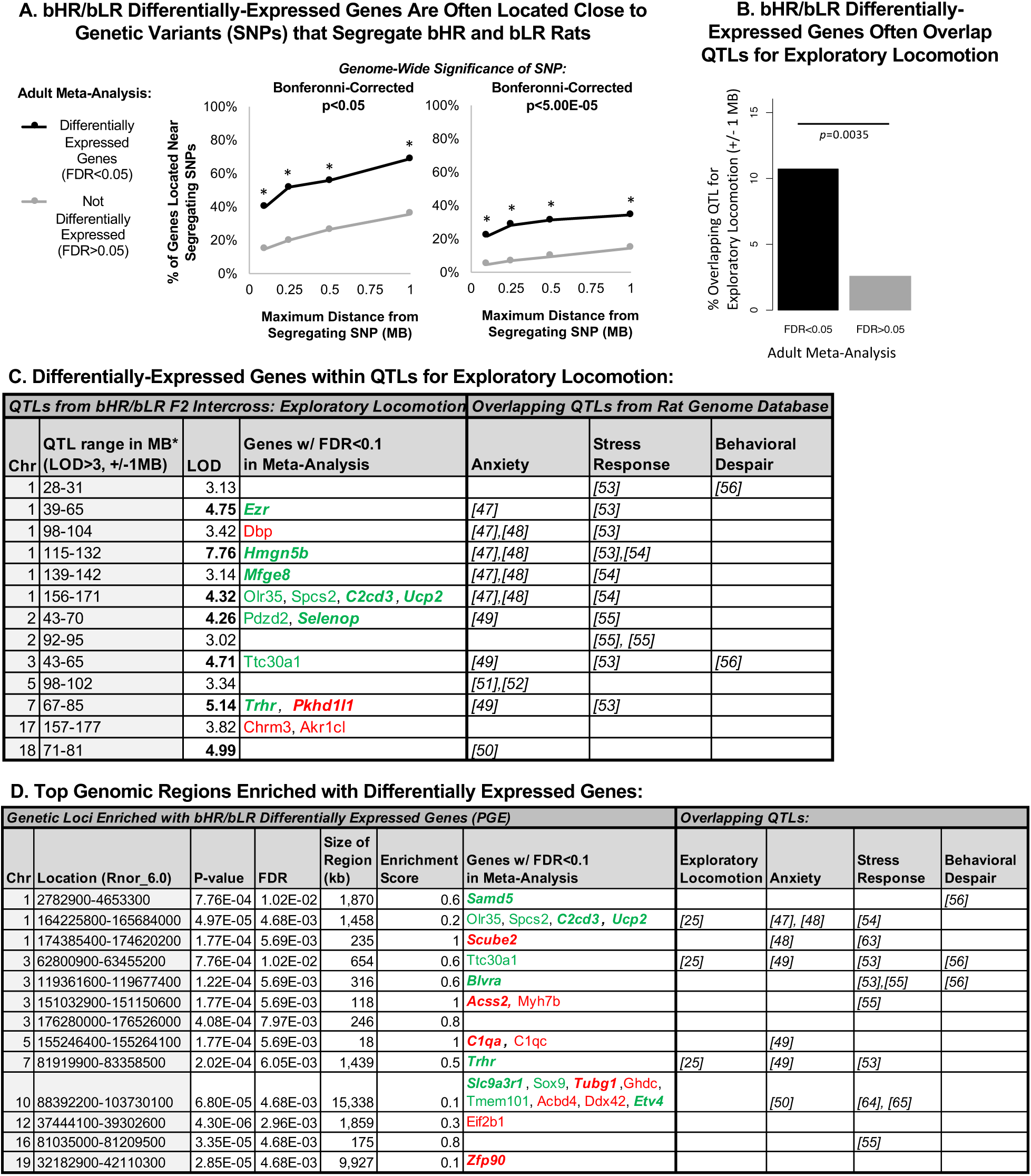
Many of the top differentially expressed genes are located near genetic variants that segregate bHR/bLR rats within quantitative trait loci (QTLs) for exploratory locomotor activity. **A.** Our concurrent genetic study used exome sequencing to identify variants (SNPs) that segregated bHR/bLR rats in our colony (25). A high percentage of bHR/bLR differentially-expressed genes (adult meta-analysis: *FDR*<0.05) were found within +/− 100 kb, 250 kb, 500 kb, or 1 MB of these segregating variants, using either traditional (Bonferonni-corrected *p*<0.05) or more stringent (Bonferonni-corrected *p*<5.00E-5) criteria to define segregation. Asterisks (*) designate enrichment (Fisher’s exact test *p*<0.0001) in comparison to the non-differentially expressed genes in our meta-analysis (*FDR*>0.05). **B.** Our concurrent genetic study used a sampling of bHR/bLR segregating variants to identify QTLs for exploratory locomotor activity in a novel field using an bHR/bLR F2 intercross (25). bHR/bLR differentially-expressed genes (adult meta-analysis: *FDR*<0.05) were 4.5x more likely to overlap (+/− 1MB) QTLs for exploratory locomotion than other genes included in our meta-analysis (Fisher’s exact test: *p*=0.0035). **C.** A table illustrating the top genes from our meta-analyses (*FDR*<0.1, bold+italic=*FDR*<0.05) that overlap (+/-1 MB) significant (LOD>4) and putative (LOD>3) QTLs for exploratory locomotion identified by our concurrent genetic study (25). Also depicted is overlap with QTLs identified in the Rat Genome Database (32) for the following behaviors relevant to the bHR/bLR phenotype: anxiety (47–52), stress-related responses (53–55), and behavioral despair (56). **D.** The top chromosomal loci enriched for bHR/bLR DE genes overlap previously-identified QTLs relevant to externalizing and internalizing behaviors. The top chromosomal loci enriched for bHR/bLR DE genes were identified using Positional Gene Enrichment analysis (PGE): location (full coordinates), p-value=nominal p-value, *FDR*=false detection rate, enrichment score= ratio of genes with *p*<0.01 out of all genes in the region. Also depicted is the overlap (+/-1 MB) of these enriched chromosomal loci with QTLs for exploratory locomotor activity (25), as well as with QTLs identified in the Rat Genome Database (32) for anxiety (47–50), stress-related responses (53–55,63–65), and behavioral despair (56).

### Positional Gene Enrichment Analysis Specifies Narrower Chromosomal Regions Contributing to bHR/bLR Phenotype

Positional Gene Enrichment (PGE) identified 132 chromosomal regions with a significant enrichment (FDR<0.05) of DE genes within the P14 and adult meta-analyses. We focused on the top regions (FDR<0.001; Figure 6D). We confirmed that most of these top loci (10/13) could be identified using recent RNA-Seq data (F37/F43), ruling out bias towards regions overrepresented on older microarray platforms. These 13 top loci were narrow chromosomal regions (measured in KB), but overlapped a strikingly high percentage of QTLs relevant to bHR/bLR phenotype. This overlap included three QTLs for exploratory activity (25), encompassing two DE genes associated with internalizing and externalizing behavior (Ucp2, Trhr; Fig 7&8), as well as 13% of the QTLs for anxiety (6/45, (47–50)), 21% of the QTLs for stress-related responses (8/38, (53–55,63–65)), and 23% of the QTLs for behavioral despair (3/13, (56)) in the Rat Genome Database (32). Therefore, these enriched loci could contain genetic variants contributing to internalizing/externalizing aspects of the bHR/bLR behavioral phenotype beyond exploratory locomotion.

**Figure 7.**
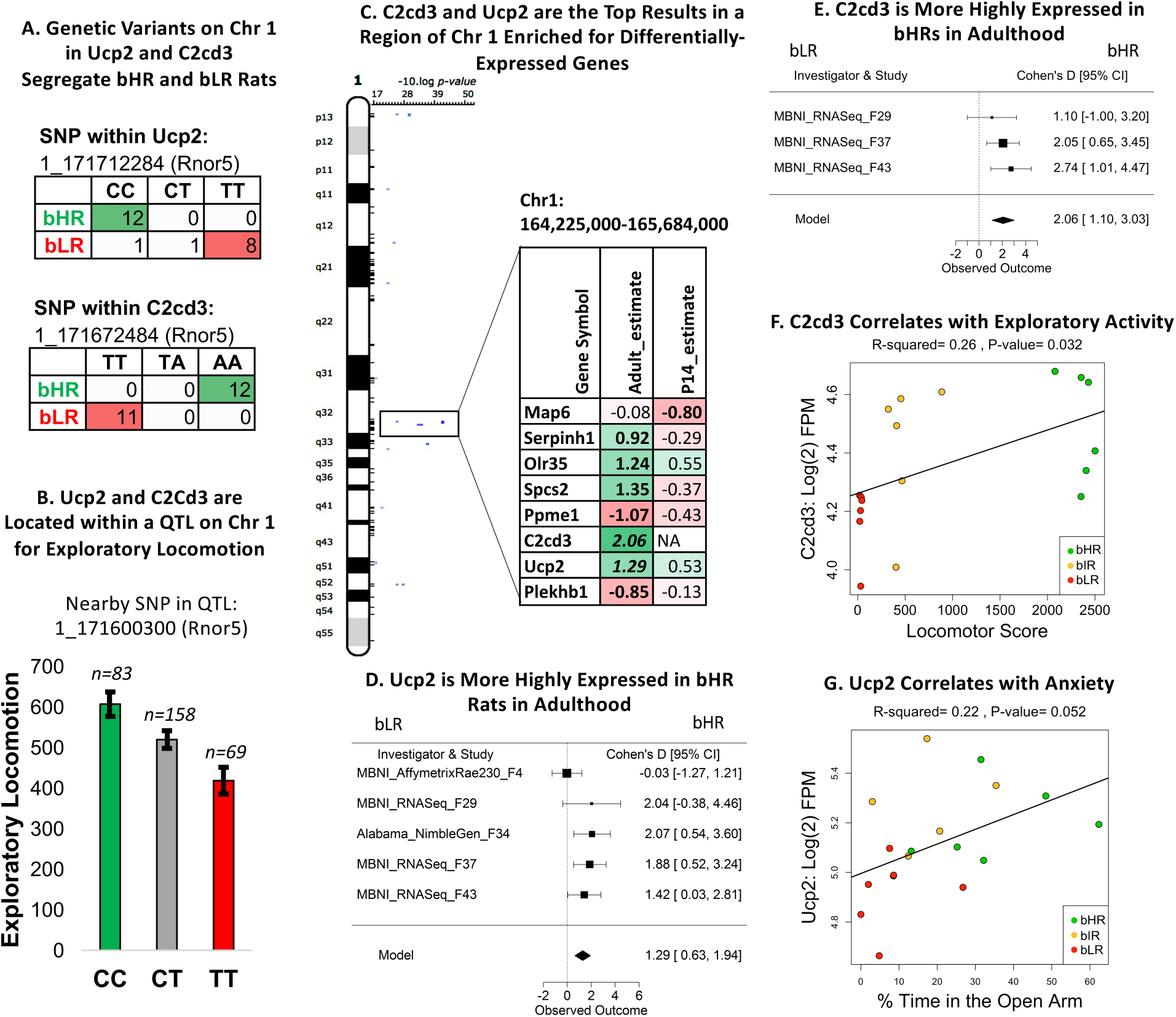
A region on chromosome 1 implicated in bHR/bLR phenotype contains two genes important for brain function and development, Uncoupling Protein 2 (Ucp2) and C2 Calcium Dependent Domain Containing 3 (C2cd3). **A)** Genetic variants on Chr 1 within Ucp2 and C2cd3 segregate bHR and bLR rats in our colony (Rnor5 coordinates, Fisher’s exact test: SNP 1_171712284: *p*=1.66E-09; SNP 1_171672484: *p*=1.27E-13). **B)** Ucp2 and C2cd3 are located on Chr 1 within a QTL for exploratory locomotor activity. An example of the correlation between genetic variation in this region and behavior is illustrated using the sequencing results from a nearby SNP and exploratory locomotor activity measured in a bHRxbLR F2 intercross (*n*=310; *adj.R^2^*=0.049, *p*=2.00E-04, *FDR*=0.0048). **C)** C2cd3 and Ucp2 were the top DE genes (*FDR*<0.05) within a segment of Chr 1 enriched for DE genes and containing a QTL for exploratory locomotor activity (25). The table illustrates the DE genes within this region: estimate=estimated effect size (green/positive=higher expression in bHRs), bold=*p*<0.05, bold+italic=*FDR*<0.05. **D)** A forest plot showing that Ucp2 had higher expression in bHRs in four out of the five adult datasets included in the adult meta-analysis (boxes=Cohen’s D from each study +/-95% confidence intervals, “Model”=estimated effect size +/-95% confidence intervals provided by the meta-analysis, effect of phenotype: *β*=1.29, *p*=1.27E-04, *FDR*=3.83E-02). This direction of effect mirrors findings in the literature showing that Ucp2 knockout mice have higher anxiety-like behavior and lower locomotor activity, as well as greater sensitivity to stress (57,58,60,61), much like our bLR rats. **E)** A forest plot showing that C2cd3 had higher expression in bHRs in three adult datasets included in the adult meta-analysis (effect of phenotype: *β*=2.06, *p*=2.84E-05, *FDR*=1.73E-02). **F)** In the behavioral data accompanying the MBNI_RNASeq_F37 dataset, C2cd3 (units: log(2) fragments per million (FPM)) showed a positive relationship with exploratory locomotor activity (*β*= 0.000109, *R^2^*=0.26, *p*=3.20E-02). **G)** In the behavioral data accompanying the MBNI_RNASeq_F37 dataset, Ucp2 showed a trend towards a positive relationship with percent of time spent in the anxiogenic open arms of the EPM (*β*=0.03, *R^2^*=0.22, *p*=5.16E-02).

**Figure 8.**
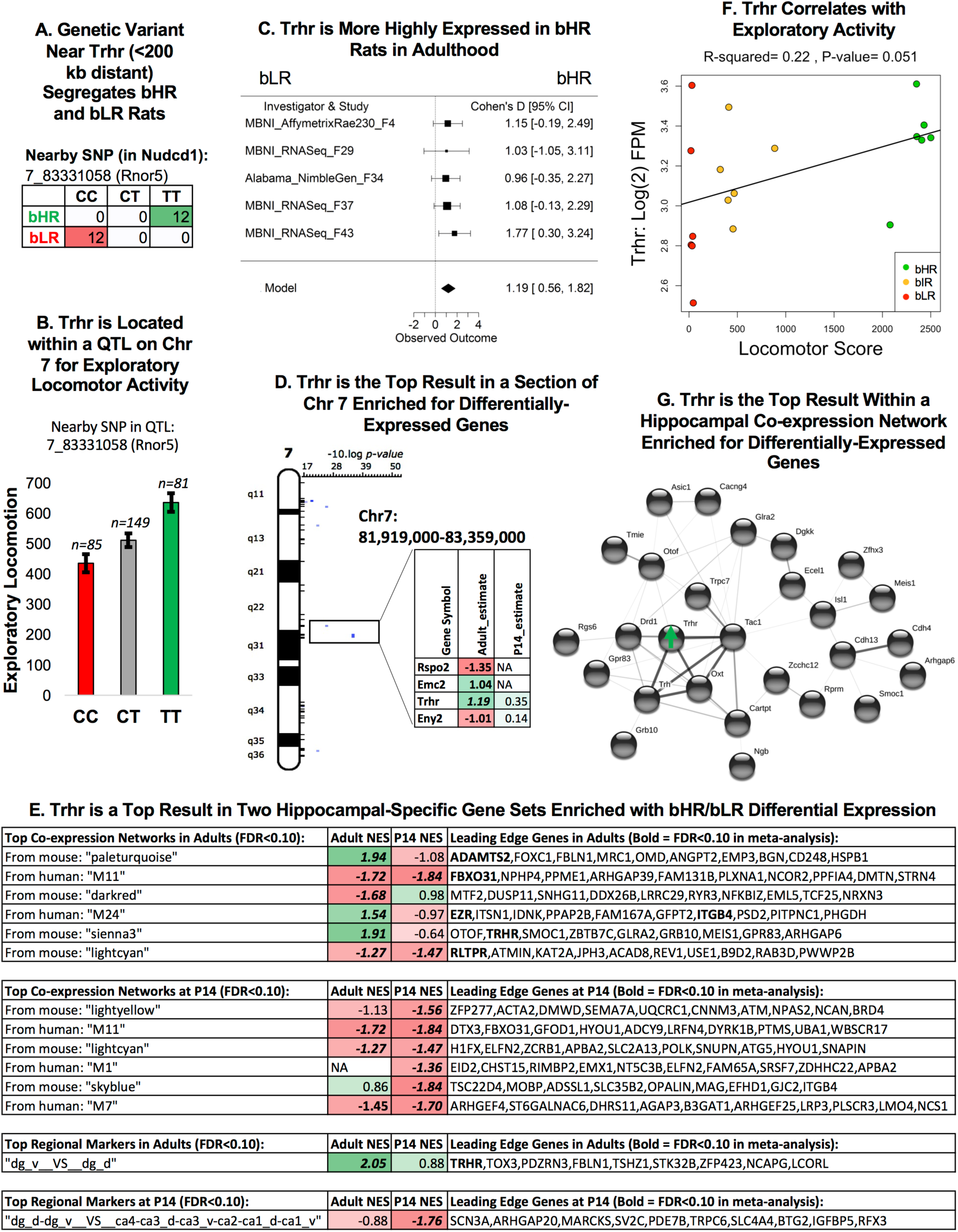
Thyrotropin releasing hormone receptor (Trhr) was the top gene within two hippocampal specific gene sets and within a region of Chromosome 7 implicated in bHR/bLR phenotype. **A)** A genetic variant on Chr 7 near Trhr (<200 kb distant) segregates bHR and bLR rats in our colony (Fisher’s exact test: *p*=6.20E-14). **B)** Trhr is located on Chr 7 within a QTL for exploratory locomotor activity. An example of the correlation between genetic variation in this region and behavior is illustrated using the sequencing results from a SNP nearby Trhr (discussed above) and exploratory locomotor activity measured in a bHRxbLR F2 intercross (*n*=315, *adj.R^2^*=0.061, *p*=5.79E-05, *FDR*=0.00178). **C)** A forest plot showing that Trhr expression was consistently elevated in bHR rats since generation F4 (boxes=Cohen’s D from each study +/-95% confidence intervals, “Model”=estimated effect size +/-95% confidence intervals provided by the meta-analysis, *β*=1.19, *p*=2.16E-04, *FDR*=4.94E-02). This direction of effect mirrors findings in the literature that show Trhr-KO mice exhibit greater anxiety and depressive-like behavior (62). **D)** Trhr was the strongest result within a segment of Chromosome 7 enriched for DE genes and overlapping a QTL for exploratory locomotor activity. The table illustrates the DE genes within this region: estimate=estimated effect size (green/positive=greater expression in bHRs), bold=*p*<0.05, bold+italic=*FDR*<0.05. **E)** Trhr was a leading gene in two of the top hippocampal-specific gene sets identified as enriched for bHR/bLR DE genes by GSEA (FDR: False Detection Rate; NES: Normalized Enrichment Score, with positive scores (green) indicating greater expression in bHRs and negative scores (red) indicating greater expression in bLRs, bold: *p*<0.05 in GSEA results, bold+italics: *FDR*<0.05 in GSEA results). The top 10 “Leading Edge” genes for each gene set are shown (bold+italics: *FDR*<0.05 in meta-analysis). These genes have large estimated effect sizes within the meta-analysis and help drive the enrichment of effects within these gene sets. Regional marker gene sets use the following abbreviations: “dg”=dentate gyrus, “v”=ventral, “d”=dorsal, “ca”=Cornu Ammonis subregion. **F)** In the behavioral data accompanying the MBNI_RNASeq_F37 dataset, Trhr (units: log(2) fragments per million (FPM)) showed a trend towards a positive relationship with exploratory locomotor activity (*β*=4.48E-04, *R^2^*= 0.22, *p*=5.08E-02). **G)** Trhr and its ligand, Thyrotropin releasing hormone (Trh), are hub genes within a hippocampal specific co-expression network that is enriched for bHR-upregulated genes. Genes within this network with known protein-protein interactions are illustrated above (STRINGdb: confidence setting=0.15 due to hippocampal co-expression already suggesting potential interaction). Many of these genes have documented associations with reward behavior.

### bHR/bLR Differential Expression is Enriched within Hippocampal Functional Pathways

#### bHR/bLR Phenotype is Associated with Proliferation and Differentiation

Eight functional ontology gene sets (out of 2,761) showed an enrichment of DE within the P14 meta-analysis (FDR<0.05), all of which were upregulated in bLRs (Fig 9A, **Table S3, Fig S6**) and predominantly related to neurogenesis, differentiation, and brain development. Within the adult meta-analysis, 2 of the 4 top gene sets enriched with DE (FDR<0.1, out of 2,761 total) were similarly related to proliferation and development, but upregulated in bHRs. This pattern was confirmed using recent RNA-Seq data (F37/F43, **Table S3**), ruling out bias towards gene families overrepresented on microarray platforms.

**Figure 9.**
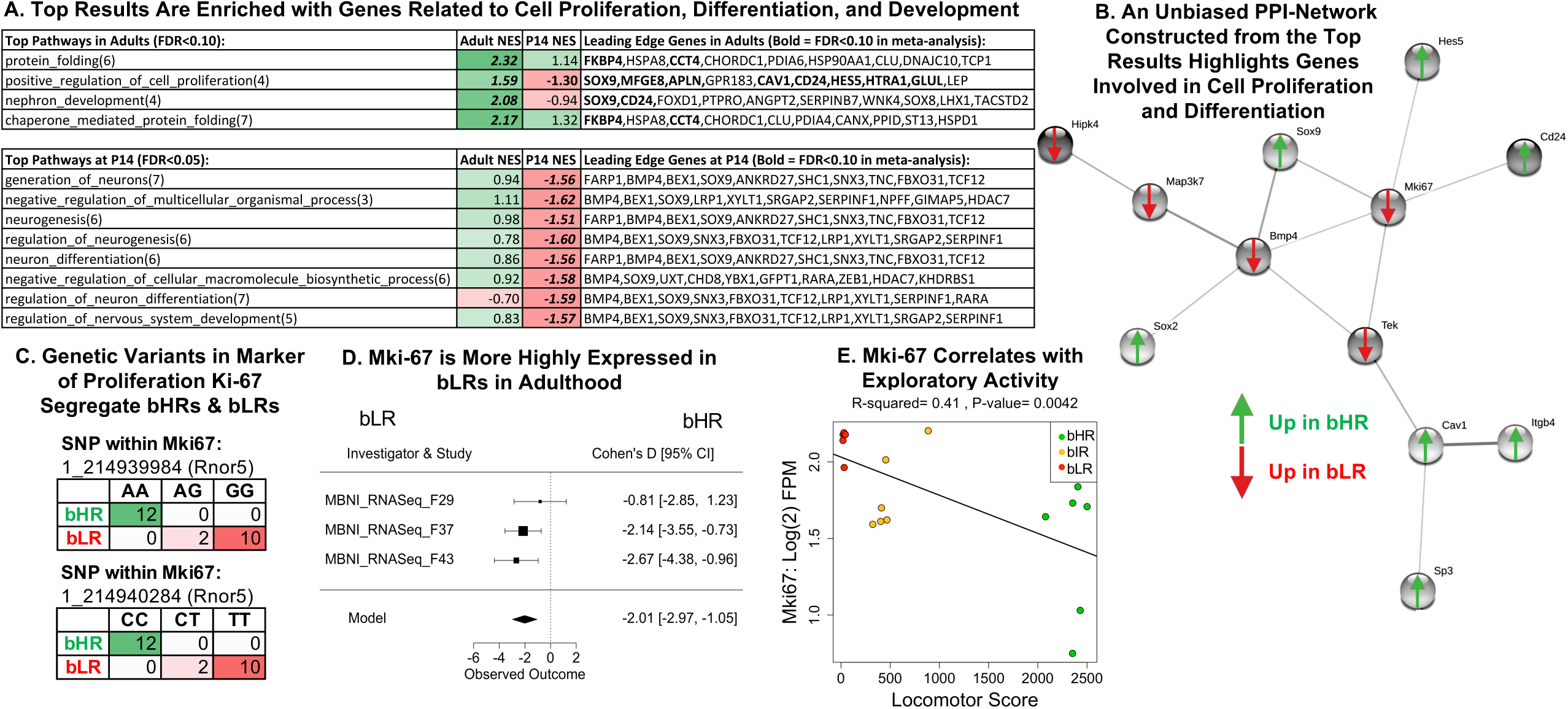
The top bHR vs. bLR DE results are enriched with genes related to cell proliferation, differentiation, and development, including the canonical Marker of Proliferation (Mki67). **A)** A table of the top functional ontology gene sets identified as enriched for bHR/bLR DE genes by GSEA (FDR: False Detection Rate; NES: Normalized Enrichment Score, with positive scores (green) indicating higher expression in bHRs and negative scores (red) indicating higher expression in bLRs, bold: *p*<0.05 in GSEA results, bold+italics: *FDR*<0.05 in GSEA results). The top 10 “Leading Edge” genes for each gene set are shown (bold+italics: *FDR*<0.05 in meta-analysis). These genes have large estimated effect sizes within the meta-analysis and help drive the enrichment of effects within these gene sets. **B)** A PPI network constructed using the top genes from the adult meta-analysis (192 genes with *FDR*<0.10, STRINGdb: confidence setting=0.40) had a dominant subnetwork that included Bmp4 and Mki67 as hub genes. Many of these genes are related to cell proliferation and differentiation within the brain. **C)** Two genetic variants on Chr 1 within Mki-67 fully segregated bHR and bLR rats in our colony (Rnor5 coordinates, Fisher’s exact test: SNP 1_214939984: *p*=2.02E-11, SNP 1_214940284*: p*=2.02E-11). **D)** A forest plot showing that Mki67 was consistently elevated in bLR rats in adulthood (boxes=Cohen’s D from each study +/-95% confidence intervals, “Model”=estimated effect size +/-95% confidence intervals provided by the meta-analysis, adult: *β*=-2.01, *p*=4.03E-05, *FDR*=1.99E-02). **E)** Within the behavioral data accompanying the MBNI_RNASeq_F37 dataset, Mki67 (units=log(2) fragments per million (FPM)) showed a negative relationship with locomotor score (*β*=-0.000249, *R^2^*=0.41, *p*=0.0042).

A PPI network constructed using the top genes from the adult meta-analysis (FDR<0.10: 192 genes) had a dominant subnetwork highlighting many of these same genes (Fig 9B), including hubs Bmp4 (*discussed above*) and the canonical marker of proliferation Mki67 (Fig 9C-E). Literature review confirmed the PPI interactions within this subnetwork and their role in proliferation and differentiation in the brain (66–78).

#### bHR/bLR Phenotype is Associated with the Dentate Gyrus (DG)

Similarly, when performing GSEA using 69 gene sets custom-designed to reflect hippocampal-specific cell types and networks (**Table S1**), we observed an enrichment of DE related to the DG (Fig 8E, **Fig S7, Table S3**), the location of neural proliferation within the hippocampus. At P14, bLRs showed an upregulation of genes with enriched expression in the DG as compared to the Cornu Ammonis (CA) regions (40) (FDR<0.05). In adulthood, bHRs showed an upregulation of genes with enriched expression in the ventral compared to the dorsal DG (40) (FDR<0.05), including Trhr (FDR<0.05, *discussed above*).

#### bHR/bLR Phenotype is Associated with Hippocampal Co-expression Networks Related to Synaptic Signaling

Co-expression modules can capture regionally-important cell types and functions that remain undocumented in traditional ontology databases (79). We observed an enrichment of bHR/bLR effects within six previously identified hippocampal co-expression modules within the P14 meta-analysis (FDR<0.05) and five within the adult meta-analysis (FDR<0.05, Fig 8E, **Fig S7, Table S3**). Two of these modules included genes within implicated chromosomal regions.

The first was a large co-expression module (695 genes) previously identified in the mouse hippocampus ((39), “*lightcyan”*) which showed elevated expression in bLRs relative to bHRs at P14 (FDR<0.05) and adulthood (FDR<0.10). One of the top DE genes in this module, ETS Variant 4 (Etv4, FDR<0.05, bHR>bLR), is a transcription factor required for proper hippocampal dendrite development (80) located within an implicated chromosomal region (**Fig S8**). A PPI network constructed using the DE genes in this module (n=74, adult p<0.05), was enriched with genes related to cell projections, neurons, synapses, and cation binding (FDR<0.05).

The second was a small co-expression module (39 genes) previously identified in the mouse hippocampus ((39), *sienna3*), which showed elevated expression in bHRs in adulthood (FDR<0.05). The top gene in this module was Trhr (*discussed above*). A PPI network constructed using all 39 genes in this module centered on Trhr and its ligand, thyrotropin releasing hormone (Trh; Fig 8G), and included many reward-related signaling molecules, including CART prepropeptide (Cartpt (81)), Oxytocin/Neurophysin I Prepropeptide (Oxt (82)), and Dopamine Receptor D1a (Drd1a (83, 84)).

#### bHR/bLR Phenotype is Associated with Microglial and Endothelial-Specific Gene Expression

Our results suggested that bHR/bLR rats might have differences in hippocampal cell type composition. A cell type deconvolution analysis focused on well-characterized non-neuronal cell type categories revealed that bLRs had greater microglial-specific expression than bHRs at P14 and adulthood (Fig 10A,C-D). At P14, bLRs also showed greater endothelial-specific expression (Fig 10B). These effects could reflect differences in cell type composition or activation state. Notably, two of the top DE genes that were located within implicated chromosomal regions are regulators of microglial state: Milk fat globule-EGF factor 8 (Mfge8, Fig 10E-H) promotes alternative (M2) activation (85), and Complement component C1q A Chain (C1qa, Fig 10I-L) promotes classical activation (86). To interrogate less well-characterized hippocampal cell types, we compared our meta-analysis results to the new mousebrain.org database (87), and found that the top DE genes (FDR<0.10) were highly expressed in a variety of cell types, including neuronal subcategories (**Fig S9**), mirroring the diversity of hippocampal functions implicated in bHR/bLR phenotype.

**Figure 10.**
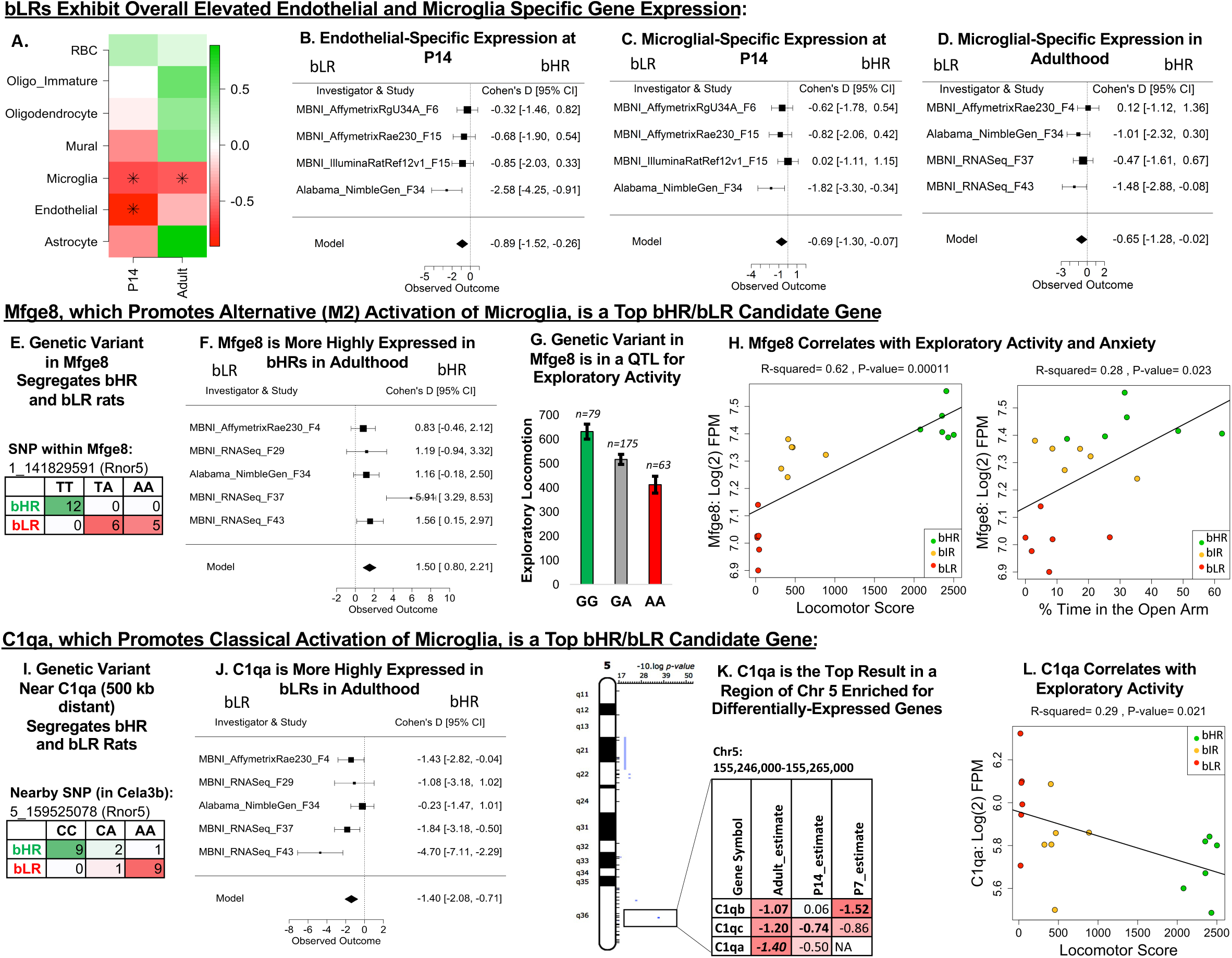
Microglial-related gene expression differentiates bHR and bLR rats. **A)** A heatmap illustrating the effect of bHR/bLR phenotype on cell type specific gene expression, which can reflect overall cell type balance (42) or activation: green=upregulated in bHRs, red=upregulated in bLRs, asterisks(*) indicate *p*<0.05. Note that only well-characterized non-neuronal cell type categories were included in this analysis. **B-D)** Forest plots (boxes=Cohen’s D from each study +/-95% confidence intervals, “Model”=estimated effect size +/-95% confidence intervals provided by the meta-analysis) showing an upregulation in bLRs of (**B**) endothelial-specific gene expression at P14 (*β*=-0.89, *p*=5.75E-03) and (**C-D**) microglial-specific gene expression **C)** at P14 (*β*=-0.69, *p*=2.90E-02) and **D)** adulthood (*β*=-0.65, *p*=4.47E-02). **E)** A genetic variant on Chr 1 in Milk fat globule-EGF factor 8 (Mfge8) segregates bHR and bLR rats in our colony (Fisher’s exact test: *p*=7.53E-08). Mfge8 promotes alternative (M2) activation of microglia. **F)** A forest plot illustrating that Mfge8 is more highly expressed in bHRs than bLRs in the adult meta-analysis in all five adult datasets (*β*=1.50, *p*=2.70E-05, *FDR*=1.73E-02). **G)** Mfge8 is located on Chr 1 within a QTL for exploratory locomotor activity. An example of the correlation between genetic variation in this region and behavior is illustrated using the sequencing results from a nearby SNP (Rnor5 coordinates 1_ 141117448) and exploratory locomotor activity measured in a bHRxbLR F2 intercross (*n*=317, *adj.R^2^*=0.061, *p*=1.81E-05, *FDR*=0.002). **H)** Within the behavioral data accompanying the MBNI_RNASeq_F37 dataset, Mfge8 (units=log(2) fragments per million (FPM)) showed a positive relationship with total locomotor score (*β*=0.000146, *R^2^*=0.62, *p*=1.10E-04) as well as the percent time in the open arms of the EPM (*β*=0.00603, *R^2^*=0.28, *p*=2.30E-02). **I)** A genetic variant on Chr 5 near Complement component C1q A Chain (500 kb distant) segregates bHR and bLR rats in our colony (Rnor5 coordinates 5_159525078, Fisher’s exact test: *p*=1.09E-07). C1qa promotes classical activation of microglia via the complement cascade. **J)** A forest plot illustrating that C1qa is more highly expressed in bLRs than bHRs in all five datasets in the adult meta-analysis (*β*=-1.40, *p*=6.57E-05, *FDR*=2.61E-02). **K)** C1qa is the top DE gene (FDR<0.05) within a segment of Chr 5 enriched for DE genes. The table illustrates the DE genes within this region: estimate=estimated effect size (red/negative=higher expression in bLRs), bold=*p*<0.05, bold+italic=*FDR*<0.05. **L)** Within the behavioral data accompanying the MBNI_RNASeq_F37 dataset, C1qa showed a negative relationship with total locomotor score (C1qa: *β*=-5.17E-04, *R^2^*=0.29, *p*=2.09E-02).

## Discussion

By selectively-breeding rats for over 16 years, we have produced a robust, genetic model of the co-occurrence of common internalizing and externalizing behaviors. Such large differences in behavior would be expected to be accompanied by similarly strong differences in gene expression in affective circuitry. By performing a formal meta-analysis across small, exploratory datasets, we provide insight into bHR/bLR differences in hippocampal gene expression across development and adulthood. Further, by cross-referencing these results with our concurrent genetic study (25), we pinpoint strong candidates for mediating the influence of selective breeding on hippocampal function and internalizing/externalizing behavior.

### Transcriptional profiling converges with genetic results to identify two strong candidates for contributing to bHR/bLR behavioral phenotype: Trhr and Ucp2

Our exome sequencing study offered a first glimpse of the genetic factors contributing to our selectively-bred phenotypes (25). However, the implicated chromosomal regions encompass several hundred genes, making their specific effects on gene expression and function difficult to predict. By cross-referencing these findings with our DE results, we discovered two strong candidates: Trhr and Ucp2. These candidates are located near bHR/bLR fully-segregating genetic variants (25) within narrow chromosomal regions highly enriched for DE genes, and overlap QTLs for exploratory activity (25), anxiety (47–49), and stress response (53, 54). Trhr was also the top gene within a bHR-upregulated gene set associated with the ventral DG (40), a region important for proliferation and emotional regulation (18), and was a hub in a bHR-upregulated hippocampal network containing reward-related signaling molecules.

Trhr and Ucp2 are both important for energy metabolism and extensively linked to internalizing and externalizing behavior (88–90). Knocking out Ucp2 produces a bLR-like phenotype: higher anxiety-like behavior, lower locomotor activity, and reduced stress resilience (57,58,60,61). Trhr is an important component of the hypothalamic-pituitary-thyroid (HPT) axis and regulates anxiety and depressive-like behavior (59,62,91). In our exome-sequencing study, the variants associated with Trhr and Ucp2 explained a moderate portion of exploratory locomotor behavior (<10%, approximately 200 locomotor counts, (25)) – a magnitude akin to the bHR/bLR difference in locomotor score present in the F1 generation. Altogether, this evidence suggests that bHR/bLR segregating genetic variants are driving DE of Ucp2 and Trhr in a manner that could meaningfully contribute to bHR/bLR differences in hippocampal function and internalizing/externalizing behavior.

### The top genes identified in the developmental meta-analyses suggest that bHR/bLR differences in hippocampal structure arise early in development: Ncan and Bmp4

The different propensity of bHR/bLR rats towards externalizing or internalizing behavior is evident at a young age (14, 15), paralleling hippocampal development (14). Our meta-analyses encompassed three postnatal ages (P7, P14, and P21) to provide insight into this neurodevelopmental trajectory. These meta-analyses depended on data from earlier generations and older transcriptional profiling platforms yet identified two strong candidates with clear associations with internalizing/externalizing behavior and hippocampal development.

The top P7 result was Ncan, exhibiting a strikingly large effect size (bHR<bLR) as early as generation F6. Ncan is located adjacent to a bHR/bLR segregating variant (25), overlapping a QTL for despair-related behavior (56). As part of the extracellular matrix, Ncan is upregulated during early brain development (92, 93) and modulates cell adhesion, migration, and growth factor binding (94). Ncan has been linked to Bipolar Disorder (92), and knocking-out Ncan enhances locomotor activity, risk-taking, hedonia, and amphetamine hypersensitivity (93). Therefore, lower levels of Ncan in early development in bHRs could promote externalizing behavior as well as divergent hippocampal development.

The top P14 result was Bmp4, which was elevated in bLRs since the earliest generations in a manner that appeared to persist into adulthood. As a regulator of development, Bmp4 is initially important for neural induction (95, 96), but later suppresses neurogenesis (97–101) and promotes other cell fates (78,95,102). Bmp4 was an important driver in bHR/bLR-enriched gene sets related to proliferation and differentiation at P14 and a hub in a related PPI network constructed from top adult DE genes. In the hippocampus, Bmp signaling promotes dorsal cell type identity and is essential for DG formation (103), matching our results indicating bLR-enrichment of gene expression related to the DG at P14 and adulthood.

Moreover, blocking Bmp signaling can produce a bHR-like phenotype, reducing anxiety, fear conditioning, and depressive-like behavior (98, 103). Therefore, Bmp4 is a strong candidate for driving long-term structural differences in the hippocampus capable of producing stable differences in temperament. However, Bmp4 was not located near a bHR/bLR segregating variant in our exome sequencing study (25), implying that impactful variation may be located in a nearby non-coding region or within a gene upstream in Bmp4’s signaling pathway.

### Functional analyses implicate hippocampal proliferation and differentiation in bHR/bLR phenotype

One of the most prominent themes in our results were functions related to cell proliferation and differentiation. Indeed, we found that Mki67 itself contained two bHR/bLR segregating genetic variants, and was more highly expressed in bLRs in adulthood (FDR<0.05), matching the upregulation in bLRs observed histologically in development (14) and maybe adulthood (9, 104). These findings confirm that the relationship between internalizing behavior and cell proliferation in our model is unlikely to be as simple as a general stunting of growth-related processes, as suggested by the neurotrophic model of stress-related mood disorders (105).

Many of our top DE genes were also important regulators of cell fate. Bmp4, SRY-box 9 (Sox9), SRY-box 2 (Sox2), Hes Family BHLH Transcription Factor 5 (Hes5), CD24 Molecule (Cd24), and TEK Receptor Tyrosine Kinase (Tek) regulate functions such as the developmental progression of neural differentiation, gliogenesis, and endothelial proliferation (66,74,75,77,106– 108). Their role in adulthood includes growth and plasticity in response to neural activity and injury (69,109–111). Therefore, these results could explain previous morphological findings indicating that cell differentiation progressed differently in the adult hippocampus in bHRs and bLRs under conditions of mild stress (9). Together, these findings raise the interesting possibility that DE within neurodevelopmental programming pathways could provide a general mechanism by which environmental stimuli, such as stress or drugs, produces divergent changes in hippocampal structure in bHR/bLR animals.

### Functional analyses implicate microglial activation in bHR/bLR phenotype

Microglial-specific gene expression was upregulated at both P14 and adulthood in bLRs, suggesting either an increased proportion of microglia cells or microglial activation. Several top candidate genes were important regulators of microglial function. bLRs had greater expression of C1qa than bHRs since the earliest generations. The C1q genes promote classical microglial activation (86) and are implicated in phagocytosis-driven synaptic pruning (86,112,113). In contrast, Mfge8 was more expressed in bHRs. Mfge8 is associated with reduced pro-inflammatory factors (114) as well as alternative (M2) microglial activation (85), playing an important role in phagocytosis (85,115,116). Ucp2, discussed previously, has anti-inflammatory function (57,58,60,117,118) and was more highly expressed in bHRs than bLRs. Both Mfge8 and Ucp2 contain bHR/bLR segregating genetic variants within probable QTLs for exploratory activity (25), suggesting that genetic variation could contribute to their DE in the hippocampus.

Together, these results seem to fit pro-inflammatory theories of internalizing disorders (119). However, we found little evidence of bHR/bLR differences in the expression of traditional inflammatory markers. Therefore, it seems more likely that bHR/bLR differences in microglial activation genes are tied to non-immune roles for microglia within the brain, including the regulation of neurogenesis, cell survival (120), and synaptic pruning (113, 121) in response to neuronal activity (113). Therefore, microglial phagocytosis could be serving as a multi-faceted tool to tailor plasticity either during development, or in response to environmental stimuli like stress or drugs of abuse.

### Conclusion and Future Directions

By comparing exome sequencing findings to hippocampal differential expression patterns during development and adulthood across many generations of selective breeding in our bHR/bLR colony, we implicate a diverse and compelling array of genes whose effects may converge to promote internalizing and externalizing behavior. Due to a dependence of our results on older platforms and exclusively male rats, we cannot claim to have identified all relevant candidates, nor have we highlighted all promising results in our text. However, we implicate two functional pathways with the capability to guide bHRs and bLRs along a divergent developmental trajectory and set the stage for a widely different reactivity to the environment. These findings will inspire new avenues of research (122–124), including cell type specific morphological analyses and the manipulation of candidate pathways to assess relevance to behavioral and neurological phenotype in our model.

## Acknowledgements

The work that was performed at the Molecular Behavioral Neuroscience Institute was supported by the following grants: Hope for Depression Research Foundation (HDRF; RGA# DTF Phase II (D) (McEwen-PI)), Pritzker Neuropsychiatric Disorders Research Consortium, National Institute on Drug Abuse Grants 5P01 DA021633-02, NIDA (5RO1 DA013386), Office of Naval Research Grant N00014-12-1-0366, Office of Naval Research (ONR) N00014-09-1-0598, Office of Naval Research (ONR) N00014-02-1-0879, NIMH Conte Center Grant #L99MH60398, and NIMH Program Project Grant #P01MH42251-01. The University of Alabama study was funded by NIMH 4R00MH085859-02 (S.M.C.). Research training for I.B. was supported by the Undergraduate Research Opportunity Program at the University of Michigan. We would also like to acknowledge everyone involved in the studies included in the meta-analysis: James Stewart, Angela Koelsch, Sharon Burke, Mary Hoverstein, Jennifer Fitzpatrick, Hui Li, Fei Li, and Amy Tang, the University of Michigan Advanced Genomics Core, and HudsonAlpha Institute for Biotechnology Genomics Services Lab. Finally, we thank everyone who has provided feedback on the manuscript: Dr. Shelly Flagel, Alek Pankonin, Pouya Mandi Gholami, Dr. Aram Parsegian, and Leah Johnson.

## Disclosures

These results were previously published in pre-print format (*bioRxiv* 774034; doi: https://doi.org/10.1101/774034). Several of the authors are members of the Pritzker Neuropsychiatric Disorders Research Consortium (MHH, RCT, FM, HA, SJW), which is supported by Pritzker Neuropsychiatric Disorders Research Fund, LLC. A shared intellectual property agreement exists between the academic and philanthropic entities of the consortium. The authors have declared that no competing interests exist.

## Notes

#### Summary of Updates

This version of the manuscript has been polished for journal submission, including massive reformatting and the addition of a co-author.

