## Supplementary material for "Genetic liability for internalizing versus externalizing behavior manifests in the developing and adult hippocampus: Insight from a meta-analysis of transcriptional profiling studies in a selectively-bred rat model": Suppl. Text and Figures

**Supplemental Methods**

**Detailed Methods for Animal Husbandry:**

***Selective Breeding:*** For the first generation, we chose 120 Sprague–Dawley rats (Charles River, Inc) to begin the bHR and bLR lines that exhibited extreme locomotion scores within a novel environment (the top and bottom 20%, respectively). Since then, we have maintained 12 breeding families in each line. For generations F1-F41, our breeding strategy amplified the differences between the two lines while minimizing inbreeding. During this time, to propagate the bHR and bLR lines we only mated the males and a females with the most extreme locomotor scores from each family. During the F30 generation, a second colony was begun at the University of Alabama-Birmingham using a selection of bHR/bLR rats from the MBNI colony. Later, at generation F42, the breeding strategy at MBNI changed to target locomotor scores closest to the designated phenotypic score (~1800 beam breaks for bHR and ~150 beam breaks for bLR) to prevent the selectively-bred lines from becoming more extreme.

***Animal Husbandry:*** On postnatal day 1 (PND1), litters were reduced to 12 pups (6 males, 6 females) and raised by their mothers within the breeder room on a 14:10 light:dark schedule (lights on 4 a.m., Standard Time). Following weaning (PND 21), the rats were housed in pairs in-house and maintained on a 12:12 light:dark schedule (lights on 6 a.m., Standard Time). Access to food and water was *ad libitum*.

**Detailed Methods for the Individual Datasets:**

**MBNI_AffymetrixRae230_F4**

This previously unpublished microarray dataset was generated at MBNI using bHR and bLR male rats from generation F4 (*n*=6 per group). The rats underwent typical locomotor testing to validate phenotype and were adults (between P102-P108) at the time of sacrifice. The rats were sacrificed via rapid decapitation followed by immediate brain extraction. The whole hippocampus, frontal cortex, and hypothalamus samples were rapidly dissected on ice, fast-frozen at –40°C, and stored at –80°C before processing. Only the hippocampal data was included in our current analysis. TRIzol reagent (Invitrogen, Calsbad, CA) was used to extract total RNA, followed by purification using RNeasy RNA purification columns (Qiagen, Valencia, CA). The quality and concentration of the RNA was determined using an Agilent bioanalyzer (Palo Alto, CA) and wavelength absorbance (260/280 nm ratio) by Nanodrop. The samples were transcriptionally profiled using Affymetrix Rat Expression Set 230 (RAE230A) microarray according to standard manufacturer’s procedures. The RAE230A arrays contained around 15,900 probe sets that primarily targeted well-annotated full-length genes. Typically, there were eleven pairs (perfect match/mismatch) of 25-mer oligonucleotide probes in each probeset. The samples were run in two batches: the first included the hippocampus and cortex samples and the second included the hypothalamus samples. Each brain region was run on a separate chip.

For our current analyses, the *ReadAffy()* function (from R package *affy* version 1.54.0; (3)) was used to read the data from the hippocampal Affymetrix .CEL files into R studio (version 1.0.153). An expression set was generated using the Robust Multi-Array Average method (RMA: (4). Using a custom .cdf (5) we summarized the probe signals into probesets for the RAE230A chip (“rae230arnentrezgcdf_19.0.0”) which mapped the probe sequences to Entrez Gene IDs (downloaded from http://nmg-r.bioinformatics.nl/NuGO_R.html on Jan 2017, release date Nov 2015). We re-annotated the data according to Official Gene Symbol using the function *org.Rn.egSYMBOL()* from the R package *org.Rn.eg.db* (version 3.4.1; (6)), and then removed rows lacking annotation (leaving 9,959 gene symbols). Quality control identified two samples with low sample-sample correlation values (around 0.66), leaving a final sample size of *n*=5 per group.

**MBNI_AffymetrixRgU34A_F6**

This dataset was obtained from a published microarray study (8) conducted at MBNI using bHR and bLR rats from the F6 generation (GEO accession #: GSE29552). Three different age groups were included: P7, P14, and P21, with *n*=6 per group (36 total). The rats were all baseline males, having undergone no behavioral testing. The rats were sacrificed via rapid decapitation, and the whole hippocampus and nucleus accumbens were immediately dissected. Only the hippocampal data was included in our current analysis. After extracting mRNA from the targeted tissue, cDNA was synthesized and run on Affymetrix Rat Genome (RG) U34A GeneChips following procedures previously described (8). This chip contained probesets targeting approximately 7,000 full-length sequences and around 1,000 expressed sequence tag clusters. Typically, there are 16 pairs (perfect match/mis match) 25-mer oligonucleotide probes in each probeset. We reanalyzed the original data (.CEL files) using methods similar to those used for the *MBNI_AffymetrixRae230_F4* dataset and a custom .cdf for the rgU34A chip (“rgu34arnentrezgcdr_19.0.0”). Following re-analysis, there were a total of 4,588 unique gene symbols present in the data. Data underwent standard quality control and no outlier samples were identified.

**MBNI_IlluminaRatRef12v1_F15**

This dataset was obtained from an unpublished microarray study conducted at MBNI using bHR and bLR rats from generation F15. The study was performed on male P14 baseline animals (*n=*6 per group). Rats were sacrificed via rapid decapitation and tissue was immediately extracted from the whole hippocampus. RNA was extracted and purified as discussed in ((8); *MBNI_AffymetrixRgU34A_F6*) and divided into two aliquots earmarked for transcriptional profiling using different microarray platforms. One aliquot of RNA was analyzed using the Illumina RatRef-12v1 Beadchip microarray following Illumina’s standard recommended procedures.

The Illumina RatRef-12v1 contains 23,375 50-mer oligonucleotide probes targeting coding transcripts. To extract the AVG_Signal for all probes for each of the samples, we used the *read.idat()* function (R package *limma* version 3.28.21; (2)). We performed a log2 transformation and then quantile normalized the data using the *normalize.quantiles()* function in the R package *preprocessCore* (version 1.34.0; (10)). The final dataset represented a total of 21,568 unique gene symbols. During quality control, a potential outlier was noted in the PCA. However, due to the strong sample-sample correlations among all samples (>0.98), removing the outlier was deemed unnecessary.

**MBNI_AffymetrixRae230_ F15**

This microarray dataset was produced as part of the same unpublished study as the *MBNI_IlluminaRatRef12v1_ F15* dataset. It used aliquots of the same RNA samples, but was run on a different platform: Affymetrix Rat Expression Set 230 A. This chip was designed with 15,900 probe sets that primarily targeted well-annotated full-length genes and some expressed sequence tags. There are typically 11 pairs (perfect match/mis match) of 25-mer oligonucleotide probes in each probeset.

We reanalyzed the original data (.CEL files) using methods similar to those used for the *MBNI_AffymetrixRae230_F4* dataset. The final dataset contained 9,958 unique gene symbols. During quality control, we found consistent irregularity associated with one subject (bLR subject X6_RN230_HC_P14_F15_L03.CEL) and it was removed prior to further analyses. Since this dataset is a technical replicate of MBNI_IlluminaRatRef12v1_F15, we considered accounting for covariation between the two sets of biological replicates in our meta-analysis using a multilevel model, but additional analyses revealed an almost perfect lack of correlation between the results produced by the two platforms (**Fig S2**). Therefore, the information that the two datasets was providing in our meta-analysis model appeared for all practical purposes independent.

**MBNI_RNASeq_F29**

This previously unpublished RNA-Seq dataset was generated at MBNI using bHR and bLR rats from the F29 generation. It included both adult (P60) and P14 rats (*n*=2 per group). These rats were all baseline males and had not undergone any behavioral testing, including standard locomotor testing. Rats were sacrificed through rapid decapitation, followed by immediate brain extraction and whole hippocampus dissection. Extraction, purification, and assessment of the quality and concentration of RNA followed similar procedures to those used for *MBNI_AffymetrixRae230_F4.*

Purified RNA was shipped to the Genomic Services Laboratory at HudsonAlpha (<https://gsl.hudsonalpha.org/index>) for short-read (50 bp length) single-end sequencing. RNA Qubit was used to assess concentration (range: 720-1120 ng/uL). The dilutions were denatured through heat application at 70˚C for two minutes and then checked for quality with a RNA Nano bioanalysis chip. Indexed non-stranded cDNA libraries were then prepared using standard Illumina procedures and a TruSeq library preparation kit (Illumina) after which they were pooled for sequencing with four samples per pool. The pooled libraries were clustered on a HiSeq flowcell (Illumina) and sequenced using a first generation HiSeq 2000 sequencer. Reads were mapped to the rat reference genome (RGSC v3.4) using Tophat2 and Bowtie2 with default parameters and a standard sequence read output. Only reads that aligned with the genome once were maintained for later differential expression analysis. Transcript assembly was collected using Cufflinks and Cuffmerge (7), and bHR vs. bLR differential expression was assessed for each age group (P14 and adult) using CuffDiff (7).

When performing our meta-analysis, we did not have access to original gene level summary data per subject at the time of the analysis, and therefore performed no further quality control or re-analysis procedures besides averaging across *t*-test statistics by gene symbol to eliminate over-representation (final unique gene symbol count in each dataset: 16,237 for adults and 16,342 for P14).

**Alabama_NimbleGen_F34**

This dataset was obtained from a published microarray study conducted at University of Alabama-Birmingham within the laboratory of a former MBNI investigator ((9); GEO accession #: GSE88874). The tissue in this study originated from the F4 generation of a new colony of selectively bred bHR/bLR rats. This new colony was created by selectively breeding F30 rats shipped from the MBNI colony, therefore we refer to their generation as F34 (F30+F4). This study included bHR and bLR baseline male rats from four different age groups: P7, P14, P21, and adult (*n*=5 per group, or 40 total). Rats were sacrificed by rapid decapitation and brains were flash frozen in isopentane and cooled by dry ice to -30˚C before being stored at -80˚C. Tissue extraction was conducted by first sectioning the brains in a cryostat to alternating sections of 20 and 300 µm, after which the 20 um sections were histologically stained to provide guidance. Tissue punches from the 300 um sections were used to extract samples from the dorsal hippocampus, amygdala, and prefrontal cortex (details in (9)). Only the hippocampal data was used in our meta-analysis.

Transcriptional profiling was conducted using NimbleGen Rat Gene Expression 12x135 microarray targeting 26,419 target genes using five 60mer oligonucleotide probes/target (135,000 probes total). Since Nimblegen uses proprietary software, we were unable to reprocess the raw microarray data and instead depended on the normalized, gene-summary data from the original study (provided on the GEO database). That said, the original data preprocessing (9) followed a pipeline very similar to our own, including RMA and quantile normalization. We performed a log2 transformation and then used the R package *org.Rn.eg.db* (version 3.4.1; (6)) to translate the original annotation (Refseq accession number) to gene symbol. After annotation, the data represented 10,674 unique gene symbols. Quality control procedures were similar to those described earlier. Both PCA and the sample-sample correlations revealed an outlier (a P21 bLR) which was removed from our analysis.

**MBNI_RNASeq_F37**

This previously unpublished RNA-Seq dataset was generated at MBNI using baseline bHR and bLR male adults from generation F37 (*n*=6 per group). Unlike the other datasets in our study, this one also included animals that showed an intermediate locomotor response to a novel field (Intermediate Responder (bIR) rats), which were obtained by cross-breeding bHR and bLR rats. We did not include the bIR data in the meta-analyses, but incorporated it later in follow-up behavioral analyses. After locomotor and anxiety testing (described in the main text), the rats were sacrificed in adulthood (bHR/bLR=P160-P167, bIR=P126-134) by rapid decapitation and the whole hippocampus was extracted on ice, rapidly frozen, and stored at -80 degrees C. Nucleotides were extracted using Qiagen AllPrep DNA RNA miRNA Universal Kit 50.

Extracted RNA was evaluated for total concentration and quality using a Nanodrop spectrophotometer (concentration range 285-432 ng/ul, 260/280 ratio range 1.61-1.80) and then sent to the University of Michigan DNA Sequencing Core (<https://seqcore.brcf.med.umich.edu)>. Before additional processing, the RNA was re-assessed for quality using the TapeStation automated sample processing system (Agilent, Santa Clara, CA) and only samples with RNA integrity numbers (RINs) of >8 were included in the analysis. The cDNA library was constructed using 0.1-3ug of total RNA and the Illumina TruSeq Stranded mRNA Library Preparation kit (Catalog #s RS-122-2101, RS-122-2102) (Illumina, San Diego, CA).

The final cDNA libraries were checked for quality once again by TapeStation (Agilent) as well as qPCR through the use of Kapa’s library quantification kit for Illumina Sequencing platforms (catalog # KK4835, Kapa Biosystems,Wilmington MA). The samples were clustered on a cBot automated cluster generation system (Illumina) for clonal amplification. The samples were then hybridized to the slide (“flow cell”) of a HiSeq 2000 (Illumina) with 6.66 samples per lane and underwent a 100 cycle paired end run in High Output mode using version 3 reagents. Following sequencing and demultiplexing, the RNA-Seq reads were aligned to the rat genome (Rnor_6.0) using the SubRead aligner (1) using default parameters with the exception of indel detection (maximum length of indel that could be detected=0). The featureCounts program (1) then generated the gene-level RNA-Seq count summaries for each sample based on ENSEMBL annotation (Ensembl v.81). This count summary dataset was then filtered to exclude rows of data from genes that did not meet a minimum threshold of 4 samples with greater than or equal to 10 counts. Rows that lacked official gene symbol annotation were also excluded.

Our current analysis used the log2 fragments per million gene-level summary output for each sample provided by the *voom()* function (R package *limma*; (2)). Quality control included 1) visualization of the overall log (base2) transformed transcript expression across all subjects via boxplot, 2) examination of the overall reads (mean and standard deviation) per subject, 3) visualization of a subject/subject correlation matrix to identify particularly atypical samples (R<.95). No outlier samples were identified.

**MBNI_RNASeq_F43**

This previously unpublished RNA-Seq dataset was generated at MBNI using adult bHR and bLR male rats from generation F43. These rats were not truly baseline, as the original study compared repeated injections of either antidepressant medication (fluoxetine or desipramine) or vehicle (VEH=1:1 saline and water) solution in bHR and bLR rats (14 days of intraperitoneal injections - P78-P92, 1 per day) under either standard laboratory housing conditions or chronic variable stress. In our meta-analysis, we included only the five male bHR VEH rats and five male bLR VEH rats housed in standard conditions. These rats did not undergo our standard locomotor testing in adulthood, but on the last day of injections they underwent social interaction testing after 15 minutes of exposure to the anxiogenic open arms of the EPM (described in main text). One hour after undergoing testing on the EPM, the animals were sacrificed by rapid decapitation and the brains were flash frozen in isopentane cooled on dry ice. The whole hippocampus was later dissected from each brain and immediately put in TRIzol™. RNA was extracted using the Zymo RNA isolation Kit and shipped to the University of Michigan DNA Sequencing Core for non-stranded short read (50-bp length) single-end sequencing using a procedure almost identical to that reported above for the *MBNI_RNASeq_F37* study.

The preprocessing procedures for the data closely follow those employed for the *MBNI_RNASeq_F37* dataset, including procedures for alignment to the reference genome (Rnor_6.0), the production of gene-level RNA-seq counts summaries for each sample using ENSEMBL annotation (Ensembl v.85), transformation into log2 fragments per million, and quality control. The read count dataset produced by featureCounts was filtered to exclude rows of data from genes that did not meet a minimum threshold of 5 samples with greater than or equal to 16 counts. After annotation, there were 16,393 unique gene symbols represented in the data. No outlier samples were identified.

**Detailed Methods for qPCR Validation:**

***Animal husbandry and tissue collection:*** Using selectively-bred rats from later generations (F51, F55), 6 bHR and 6 bLR males were sacrificed at ages P14 and P90 (**Fig S1**). For the P14 collection, the rats were sacrificed within 3-5 min of separation from the dam. The rats designated for the P90 collection were weaned, housed, and loco-tested as previously described. Sacrifice was performed via rapid decapitation without anesthesia. Brains were immediately hemisected and flash frozen by submersion in -30°C 2-methylbutane.

***RNA extraction and cDNA synthesis:*** Brains were stored at -80°C for fewer than 6 months before processing. Hippocampus was dissected from one hemisphere and homogenized using a QIAshredder kit (Qiagen #79654), and RNA was extracted using an RNeasy Mini Kit (Qiagen #74104). cDNA was synthesized using a 20 μL reaction containing 400 ng of RNA template (iScript cDNA Synthesis Kit, Biorad#1708891).

***qPCR:*** The primers were custom-designed to target Bmp4 (ACC# NM_012827.2; forward primer: 5’-CCCTGGTCAACTCCGTTAAT-3’, start = 1214; reverse primer: 5’-AACACCACCTTGTCGTACTC-3’, start = 1319) and the reference gene Gapdh (ACC# NM_017008.4; forward primer: 5’- GTTTGTGATGGGTGTGAACC-3’, start = 459; reverse primer: 5’-TCTTCTGAGTGGCAGTGATG-3’, start = 628). Calibration curves for the Bmp4 and Gapdh primers were constructed using a standard dilution analysis of stock cDNA that had been previously synthesized from a mixture of adult and P14 bHR and bLR hippocampi (H20, 0.1 uL, 0.5 uL, 1 uL, 2 uL) in triplicate using iQ^TM^ SYBR® Green Supermix. The bLR/bHR samples were then analyzed using a similar procedure in triplicate, with the samples from each time point processed within a separate batch.

***Data analysis:*** The calibration curves revealed efficiencies close to 1 (Bmp4: R^2^=0.98, Gapdh: R^2^=0.99, **Fig S1**), therefore the sample data for each time point was analyzed using the traditional Livak method (11, 12). After averaging the triplicate quantification cycle (C_q_) values for each sample for each probe, the data was normalized by subtracting the C_q_ for the reference gene (Gapdh) from the target (Bmp4) for each sample (ΔC_q_). Group differences in ΔC_q_ were assessed using Welch’s two sample t-test (12). Group differences in the Cq values for the reference gene (Gapdh) were also examined as a control (12). To provide an intuitive comparison to the microarray and RNA-Seq data, the data were plotted as log(2) fold change values (-ΔΔC_q_), by subtracting the average ΔC_q_ for the bLRs from all ΔC_q_ values, and then multiplying by -1.

**Additional Methods for Collective Analysis Procedures:**

***Examining the Relationship Between Gene Expression and Behavior:*** For the two adult datasets that contained associated behavioral data we used simple bivariate linear models to explore the relationship between gene expression and behavior: total locomotor score, percent time in the open arm of the EPM, and percentage of time spent interacting socially following exposure to a single mild stressor. A general effect of phenotype on these variables was evaluated using ANOVA (Type 3). We also examined the number of fecal boluses excreted during the EPM test served as a measure of anxiety independent from spontaneous exploration (13), but these results are not discussed due to the general redundancy of the findings with more traditional EPM measures.

***Effect Size Calculation:*** We calculated the effect size (Cohen’s d and variance of d) for the effect of bHR/bLR phenotype on log(2) gene expression (hybridization signal or FPM) within each age group (R package *compute.es* (14)). The raw data for the small MBNI_RNASeq_F29 dataset were inaccessible at the time of analysis, so we re-derived effect sizes from previous *CuffDiff* t-statistic output (7), averaging the few results representing duplicated gene symbols. To visually compare effect sizes across datasets we created forest plots using *forest.rma()* in the package *metafor* (15)).

***Comparison with Genetic Results:*** The original bHR vs. bLR exome sequencing results targeted 129,237 genetic variants (single nucleotide polymorphisms or SNPs) in 12 bHR and 12 bLR F37 rats, identified by Rnor5 coordinates (16). To identify QTLs for exploratory locomotor behavior, 416 of the SNPs that segregated bHR and bLR rats were then targeted in an F2 bHRxbLR intercross (*n*=314), and then QTL peaks were identified using the Haley-Knott (H-K) regression method with 1 cM resolution (16). To compare the exome sequencing results with our differential expression findings from the adult meta-analysis, we accessed the coordinates for all genes included in the adult meta-analysis (Rnor_6 annotation from the R package *org.Rn.eg.db* (6)) and used the NCBI Genome Remapping Service (<https://www.ncbi.nlm.nih.gov/genome/tools/remap>, accessed 8/8/2019 using the first 100 bp of each gene and default parameters).

To determine whether there was an enrichment of differential expression nearby segregating variants, we filtered the exome sequencing results to only include results from genes located on the 20 autosomes and 2 sex chromosomes (Rnor_5). We then examined the percentage of differentially expressed genes (adult meta-analysis FDR<0.05) found nearby segregating variants as defined using either standard (Bonferonni-corrected p<0.05) or more stringent criteria (Bonferonni-corrected p<5.00E-05). The distances examined included within +/-100 kb, 250 kb, 500 kb, or 1 MB of the segregating variants. Enrichment was defined using Fisher’s exact test to evaluate the cross table for differential expression vs. presence of a nearby segregating variant for each of the 8 permutations of stringency & distance.

To determine whether there was an enrichment of differential expression within QTLs for exploratory locomotor behavior, for each gene symbol we extracted the logarithm of the odds (LOD) score from the QTL analysis for the 1 MB bin that encompassed the gene and the two adjacent (+/- 1 MB) bins (Rnor_5), and then calculated the maximum. Enrichment was defined using Fisher’s exact test to evaluate the cross table for differential expression (adult meta-analysis FDR<0.05) vs. location within 1 MB of a QTL peak (using the cut-off for genome-wide significance in (16): LOD>4).

The results from these genetic analyses were compared to QTLs identified in the Rat Genome Database ((18): accessed 08/08/2019 with key words “Anxiety”, “Stress”, and “Despair”, and Rnor6 chromosomal coordinates) using Rnor_6 annotation. The top enriched chromosomal loci (p<0.001, FDR<0.01) identified by Positional Gene Enrichment Analysis (20) were also converted to with Rnor_6 annotation for reporting and comparison with our genetic findings by cross-referencing the encompassed genes with available coordinates (<https://www.ncbi.nlm.nih.gov/genome/gdv/?org=rattus-norvegicus>).

***Gene Set Enrichment Analysis (GSEA):*** The gene lists were ranked according to the beta coefficient values (β) representing the magnitude and direction of effect estimated by the meta-analysis. We performed GSEA using the *fgsea()* function (R package *fgsea* (43); settings: 100,000 permutations, maxSize=1000) and gene set matrix files (.gmt) containing either standard gene ontology for rats (go2msig.org; (37): *“Rattus_norvegicus_GSEA_GO_sets_all_symbols_highquality_April_2015.gmt”* downloaded 6/2017), or customized hippocampal-specific gene sets **(Table S1)** from: *1)* human hippocampal co-expression modules from post-mortem and freshly-resected tissue (45); *2)* mouse hippocampal co-expression modules from behaviorally-profiled animals in the hybrid mouse diversity panel (46); *3)* sets of genes with expression specific to hippocampal neuronal subtypes or subregions (*Hipposeq:* (47)).

***Expression Within Particular Cell Types:*** To add additional annotation for the top differentially expressed genes (FDR<0.10), we used the new mousebrain.org database (17), which contains information about basal gene expression within a wide variety of cell types in the nervous system as identified using a large (n>500,000) mouse single-cell RNA-Seq dataset. To perform this analysis, we only considered the 81 cell types (clusters) that were likely to be present in our whole hippocampal dissections (as defined by a tissue source, probable location, and/or region of expression that included the hippocampus). These 81 cell types were subcategories within six general cellular classes (oligodendrocytes, vascular, immune, astrocytes, ependymal, neurons). We determined whether each of our top differentially-expressed genes were reliably expressed in a particular hippocampal cell type (trinarization score for cluster >0.95) and then assessed the enrichment of that expression as compared to cell types within the other classes of cells.

**Supplemental Results**

**Additional Co-expression Networks Enriched with bHR/bLR Differentially Expressed Genes**

Co-expression modules can capture regionally-important cell types and functions that remain undocumented in traditional ontology databases (19). We observed an enrichment of bHR/bLR effects within six hippocampal co-expression modules within the P14 meta-analysis (FDR<0.05) and within five co-expression modules within the adult meta-analysis (FDR<0.05, **Fig S6**). For the sake of conciseness, within the main text we only highlighted the results from two of these co-expression modules (*lightcyan, sienna3*)– additional results are described here.

*M11*, a medium-sized co-expression module (149 genes) present in both healthy and diseased hippocampi in humans, as well as in mouse hippocampi (21), was interesting because it showed an enrichment of bHR/bLR effects both within the P14 and adult meta-analyses (FDR<0.10) with expression generally elevated in bLRs compared to bHRs (NES<0). However, during further investigation, we could not assign a clear function to the *M11* module – it contained no genes exhibiting robust bHR/bLR effects (FDR<0.05), no enrichment of documented PPI or traditional functional pathways, and only two genes with differential expression in both development and adulthood (Fbox31, Arhgap39, p<0.05).

Of the remaining seven hippocampal co-expression modules that exhibited an enrichment of bHR/bLR effects in either P14 or adulthood, four seemed particularly provocative. Within the P14 meta-analysis, the *skyblue* module (top genes: Mobp and Mag) showed higher expression in bLRs compared to bHRs (NES<0). This module was highly-enriched for oligodendrocyte-specific gene expression, as indicated by overlap with the *BrainInABlender* database (64%, (22)). The large module, *M1* (1,225 genes), was also upregulated in bLRs in the P14 meta-analysis. It had been previously identified in both healthy and diseased hippocampi in humans, as well as in mouse hippocampi, and showed an enrichment of pathways related to synaptic processing (21). This module was led by the Apba2 gene (*β*=-1.03, *p*=0.0015, FDR=0.58) which was recently shown to contain both a segregating variant in relationship to bHR/bLR phenotype and a strong QTL for locomotor behavior (16). This module also contained many genes with a strong relationship to bHR/bLR phenotype in adulthood (FDR<0.05, Fxyd7, Tmem2, Rltpr, Tubg1, Slc9a3r1).

Within the adult meta-analysis, the *paleturquoise* module (top gene: Adamts2) showed greater expression in bHRs (NES>0) and displayed a strong cell type association, with 34% of the genes in the module specifically expressed in either astrocytes or vasculature, as indicated by overlap with the *BrainInABlender* database (22). The *darkred* module (top gene: Dusp11), which showed greater expression in bLRs in the adult-meta-analysis, showed a nominal positive correlation with contextual fear immobility in a previous study (*p*=0.01; (23)).

**Supplemental Table Legends**

**Table S1. For the Gene Set Enrichment Analyses: a .gmt containing 69 gene sets custom-designed to reflect hippocampal-specific functions.**

**Table S2. The full meta-analysis results for each age group (adult, P7, P14, P21), as well as a meta-analysis performed using just the adult RNA-Seq data from the latest generations (F37 & F43).**

**Table S3. The full output from the Gene Set Enrichment Analyses conducted using the results from the adult and P14 meta-analyses, including a version of the adult meta-analysis conducted using just RNA-Seq data from the two latest generations (F37 and F43).**

**Supplementary Figures**

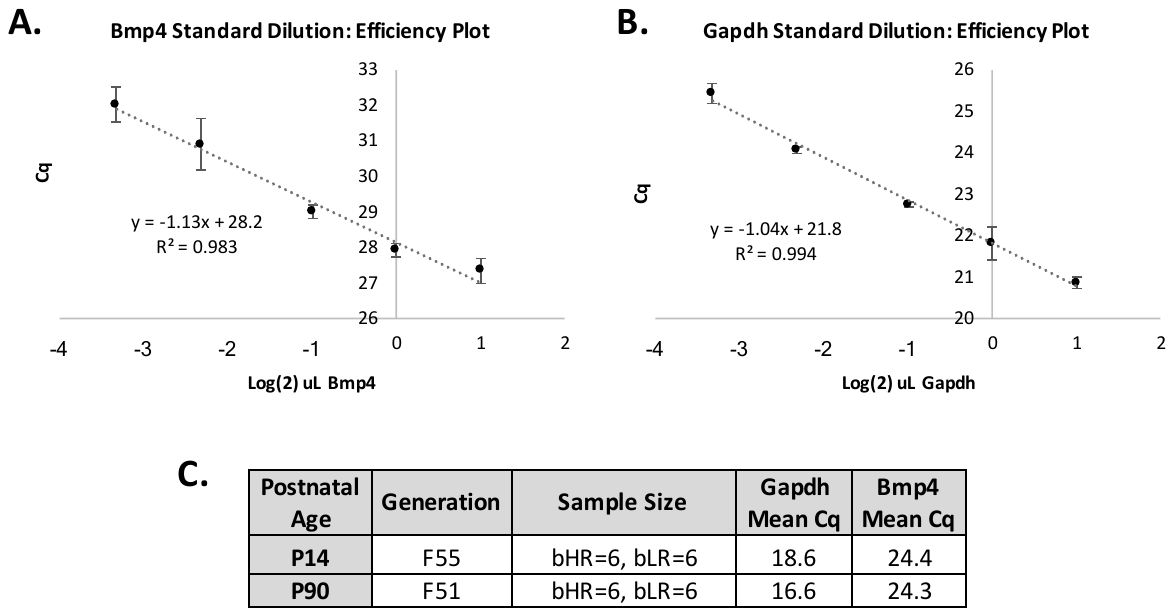

**Fig S1. Efficiency curves and methodological detail to accompany the qPCR validation results. A-B)** The calibration curves using stock cDNA revealed efficiencies close to 1 for each probe (Bmp4: *ß*= -1.13, R^2^=0.98, Gapdh: *ß*=-1.04 R^2^=0.99), implying a doubling of PCR product with each cycle. C_q_= the number of PCR cycles necessary to reach threshold. **C)** Overview of the samples used for the qPCR validation experiment. Note that the mean Cq for both the target and reference probes (Gapdh, Bmp4) was less than the Cq’s measured within our calibration experiment, indicating a greater concentration for both transcripts in our samples than in the stock cDNA. Therefore, our estimates of the concentration for both transcripts should have safely occurred during the exponential phase of the PCR procedure, prior to depletion of PCR reagents (12).

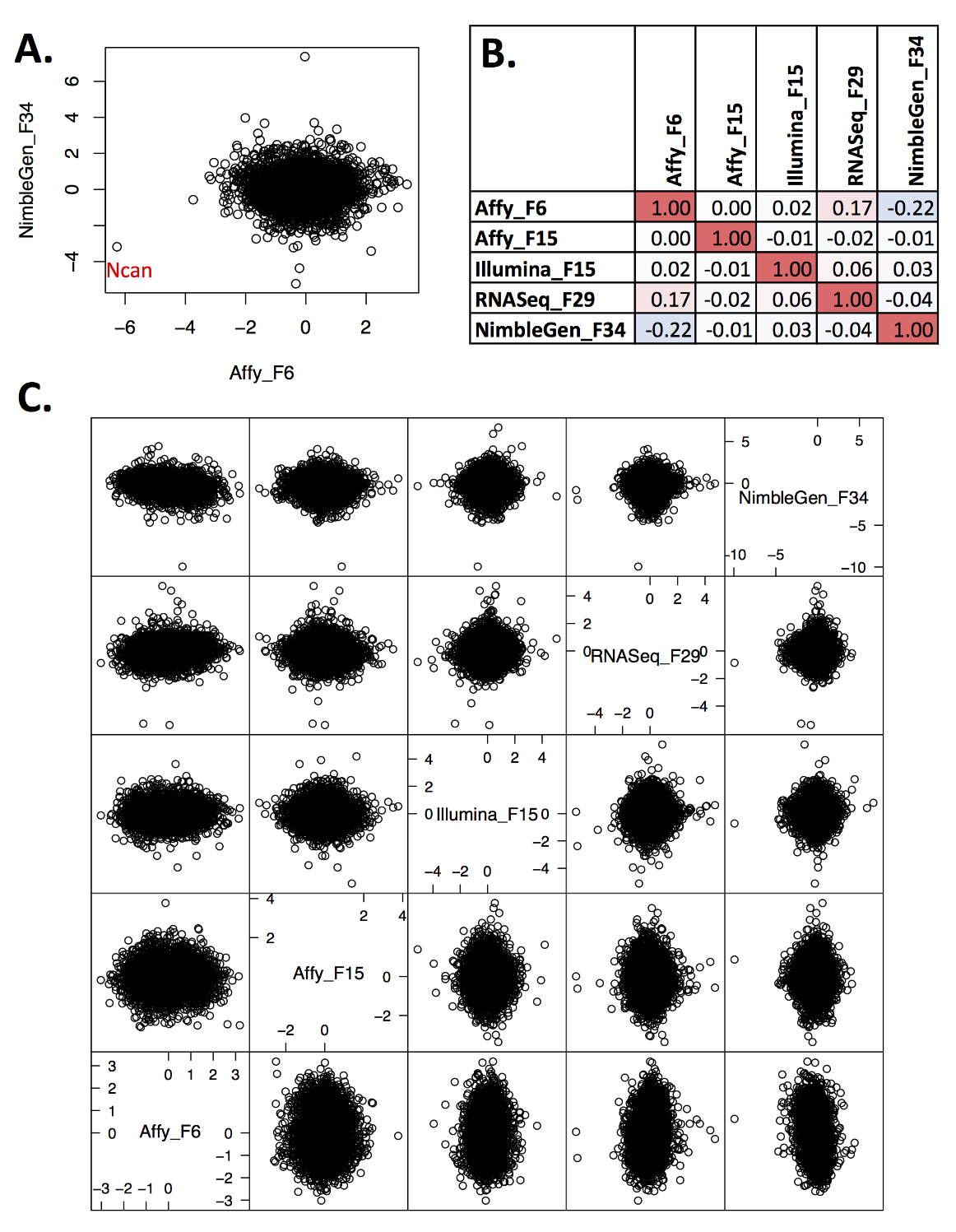

**Fig S2. The results from small (n<6) developmental bHR/bLR transcriptomic studies are noisy and produce few reliable results when analyzed individually. A)** A scatterplot illustrating the lack of correlation between the bHR vs. bLR effect sizes (Cohen’s d) for all genes included in both P7 microarray studies. Neurocan (Ncan) is the only gene showing a strong effect in both studies. **B)** The pairwise correlation coefficients between each of the bHR/bLR P14 transcriptomic studies are weak (R<0.17). The correlation coefficients were calculated using the bHR vs. bLR effect sizes for all genes present in each pair of datasets. **C)** Scatterplots illustrating the lack of correlation between the P14 transcriptomic studies. Note that the correlations are not noticeably better between studies with similar platforms or identical generations – most of the noise is likely to originate directly from the usage of small sample sizes.

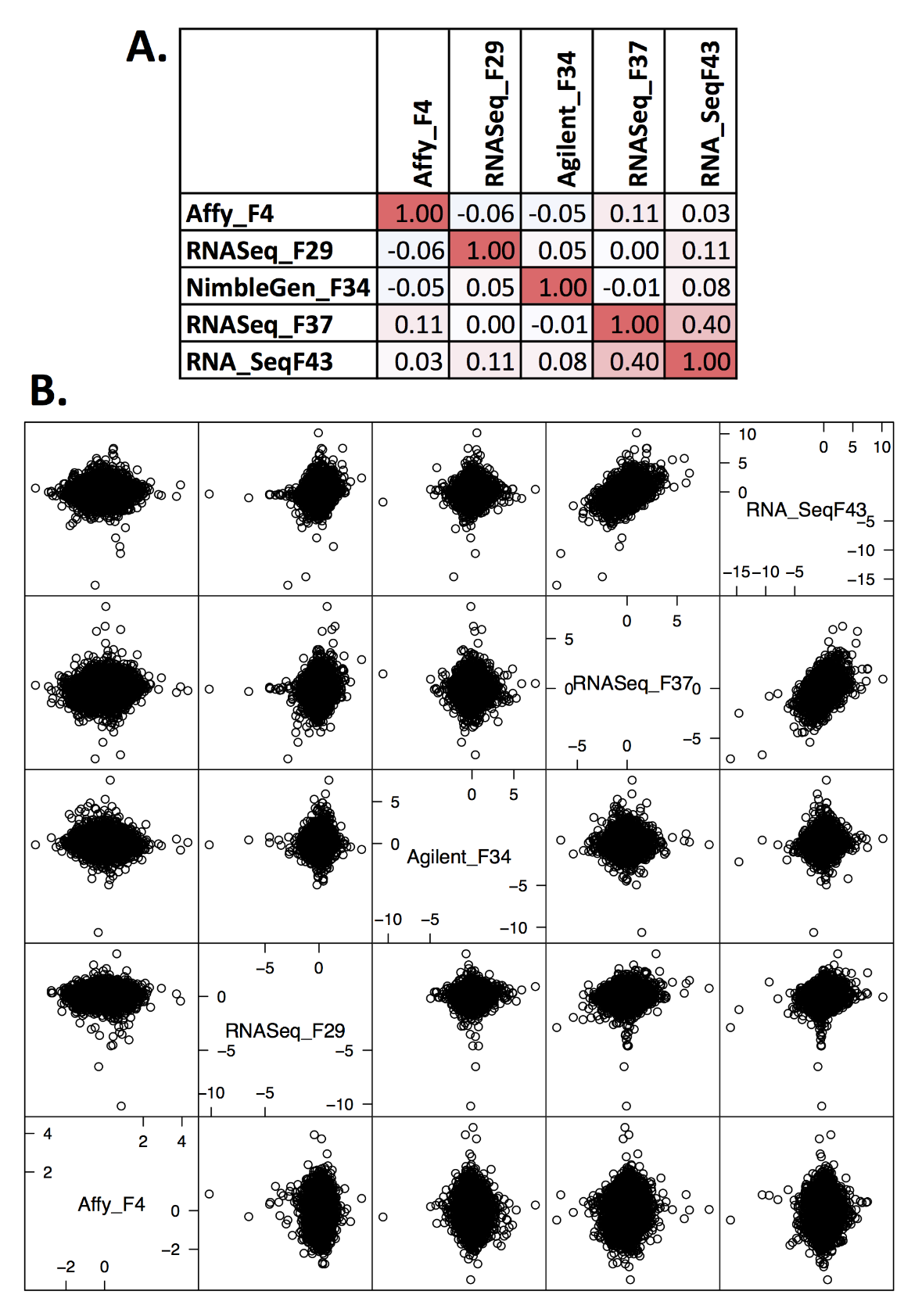

**Fig S3. The results from small (n<6) adult bHR/bLR transcriptomic studies are noisy and produce few reliable results when analyzed individually until the latest generations of selective breeding. A)** The pairwise correlation coefficients between each of the bHR/bLR adult transcriptomic studies are weak (R<0.11) until the latest generations (F37 and F43, R=0.40). The correlation coefficients were calculated using the bHR vs. bLR effect sizes (Cohen’s D) for all genes present in each pair of datasets. **B)** Scatterplots illustrating the lack of correlation between the adult transcriptomic studies until the latest generations.

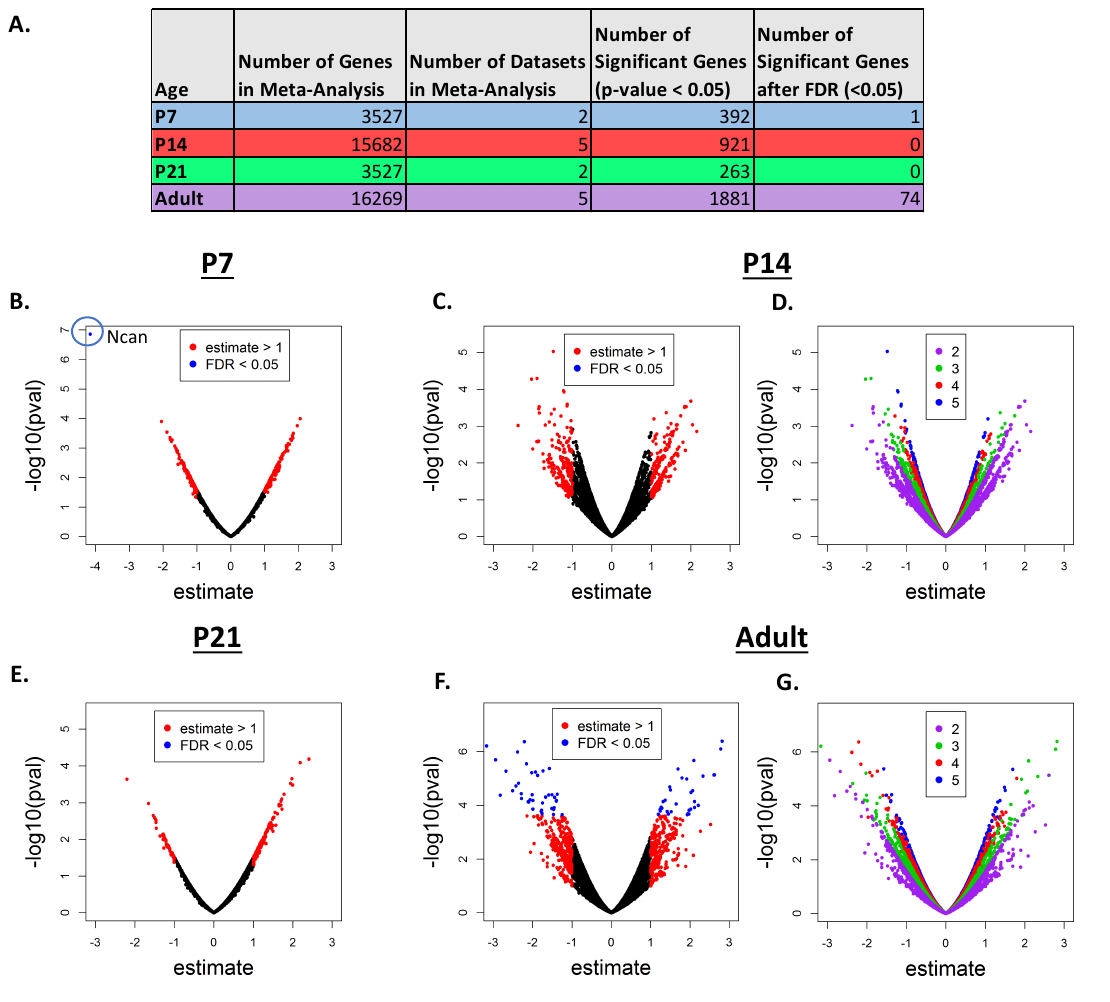

**Fig S4. Meta-analysis uncovers hippocampal gene expression related to bHR/bLR phenotype: An overview. *A)*** A table overviewing the results of the meta-analyses conducted for each of the four age groups. Genes were only included in the meta-analysis if they were represented in at least two datasets. As a result, the P14 and Adult meta-analyses contained a much larger number of genes because they contained five datasets, whereas the P7 and P21 meta-analyses each contained two datasets. The most DE genes were detected within the adult meta-analysis, which predominantly included data from later generations (F29-F43). **B-G.** Volcano plots illustrating the distribution of the results for all genes included in the meta-analysis for each age group. The x-axis is the estimated effect size for each gene (i.e., the difference in expression between bHR and bLR rats in units of standard deviation, positive=higher expression in bHRs), the y-axis is the -log(10) nominal p-value from the meta-analysis (larger=more significant). These plots are colored to either illustrate: **B-C & E-F)** the number of genes in each meta-analysis with an estimated effect size greater than one (red) or with p-values surpassing false detection correction (FDR<0.05, blue), **D&G)** The number of datasets included in the meta-analysis for each gene (for the P14 and adult meta-analyses). We found that the estimated effect sizes tended to be more extreme for genes present in fewer datasets, most likely because those genes were often only represented in the RNA-Seq datasets from later generations.

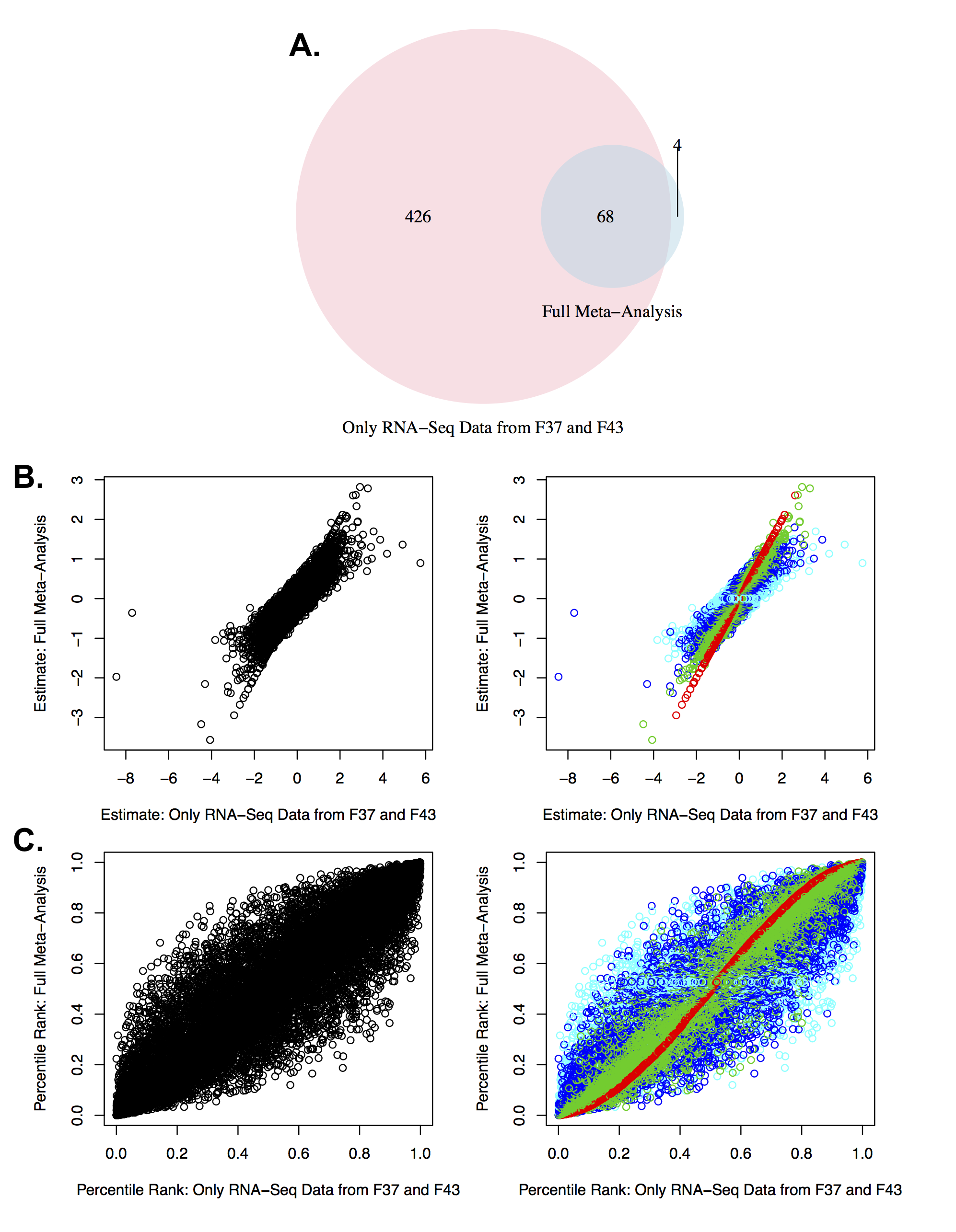

**Fig S5. A meta-analysis of the adult RNA-Seq data from only the latest generations (F37 and F43) re-identifies almost all (68/74) of the top genes discovered in the full adult meta-analysis. A)** A Venn diagram illustrating the overlap between the top genes (FDR<0.05) identified within the adult meta-analysis when using only the two datasets from the most recent generations (F37 and F43) versus all datasets available for any particular gene. There are more genes identified as related to bHR/bLR phenotype when only considering the data from the most recent generations. **B-C)** Scatterplots illustrating the strong correlation between the estimated effect sizes (B) and percentile ranks (C) produced by the two versions of the meta-analysis. The scatterplots on the right are colored to illustrate the number of datasets included in the full meta-analysis (turquoise=5, blue=4, green=3, red=2).

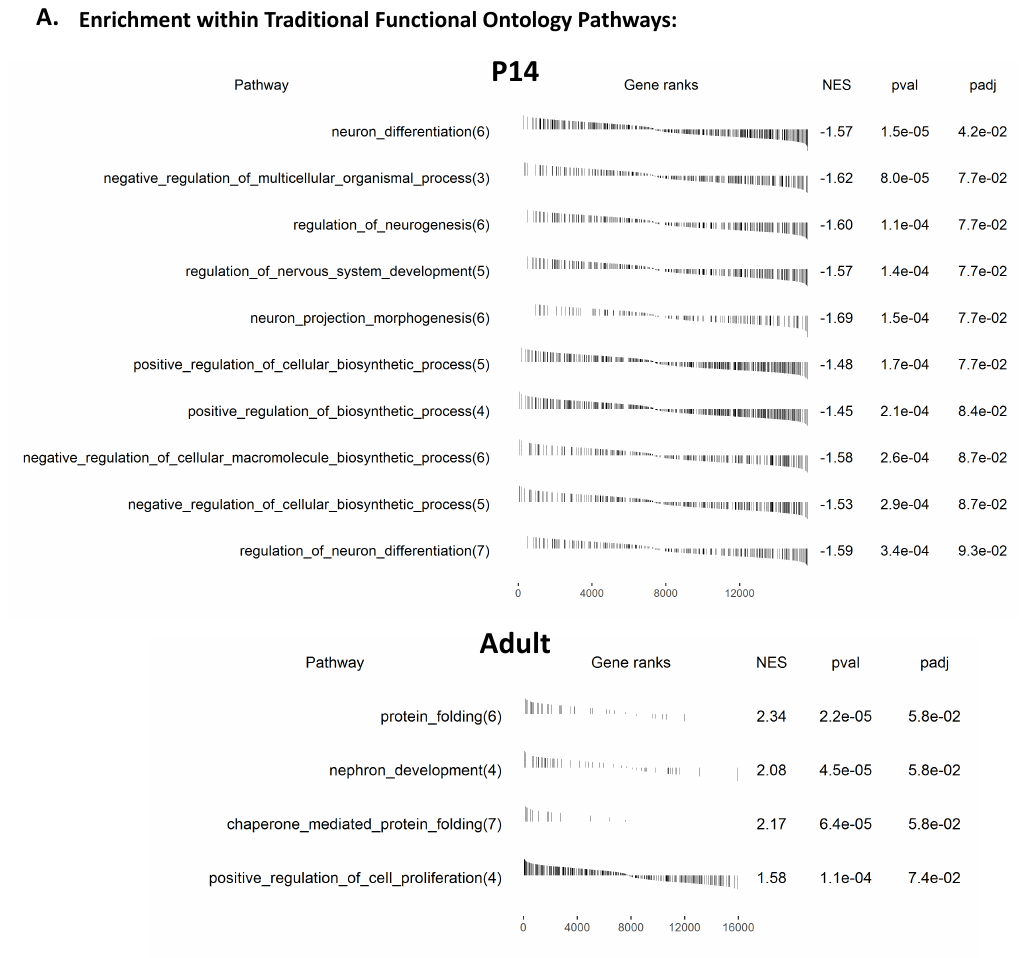

**Fig S6. The top results (FDR<0.10) from gene set enrichment analysis for the P14 and adult meta-analysis performed using a database (.gmt) of traditional functional ontology pathways.** The gene ranks were determined using the estimate from the meta-analysis (lowest rank=highly upregulated in bHRs (largest positive estimate), highest rank=highly upregulated in bLRs (largest negative estimate)) and the dark lines illustrate the rank and estimate for each gene in the respective gene set. NES=normalized enrichment score for the gene set (positive=upregulated in bHRs, negative=upregulated in bLRs), pval=nominal p-value, padj=false detection rate.

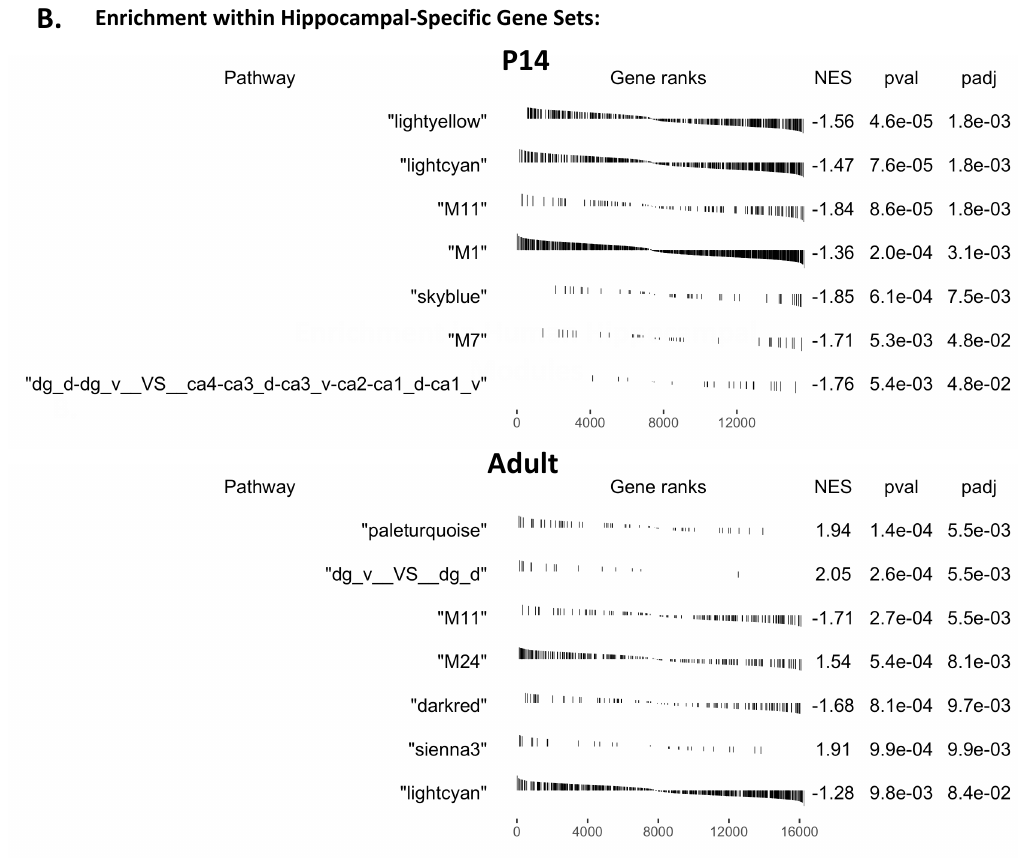

**Fig S7. The top results (FDR<0.10) from gene set enrichment analysis for the P14 and adult meta-analysis performed using a database (.gmt) of hippocampal-specific co-expression networks and gene sets*.*** Formatting is identical to Fig. S6. Gene sets named after colors represent co-expression networks previously identified in a large mouse study (23), whereas the gene sets identified with module numbers (M#) represent co-expression networks previously identified in a large human study (21). The other gene sets are derived from hippocampal regional comparisons of gene expression ((24); dg=dentate gyrus, d=dorsal, v=ventral, ca=Cornu Ammonis regions).

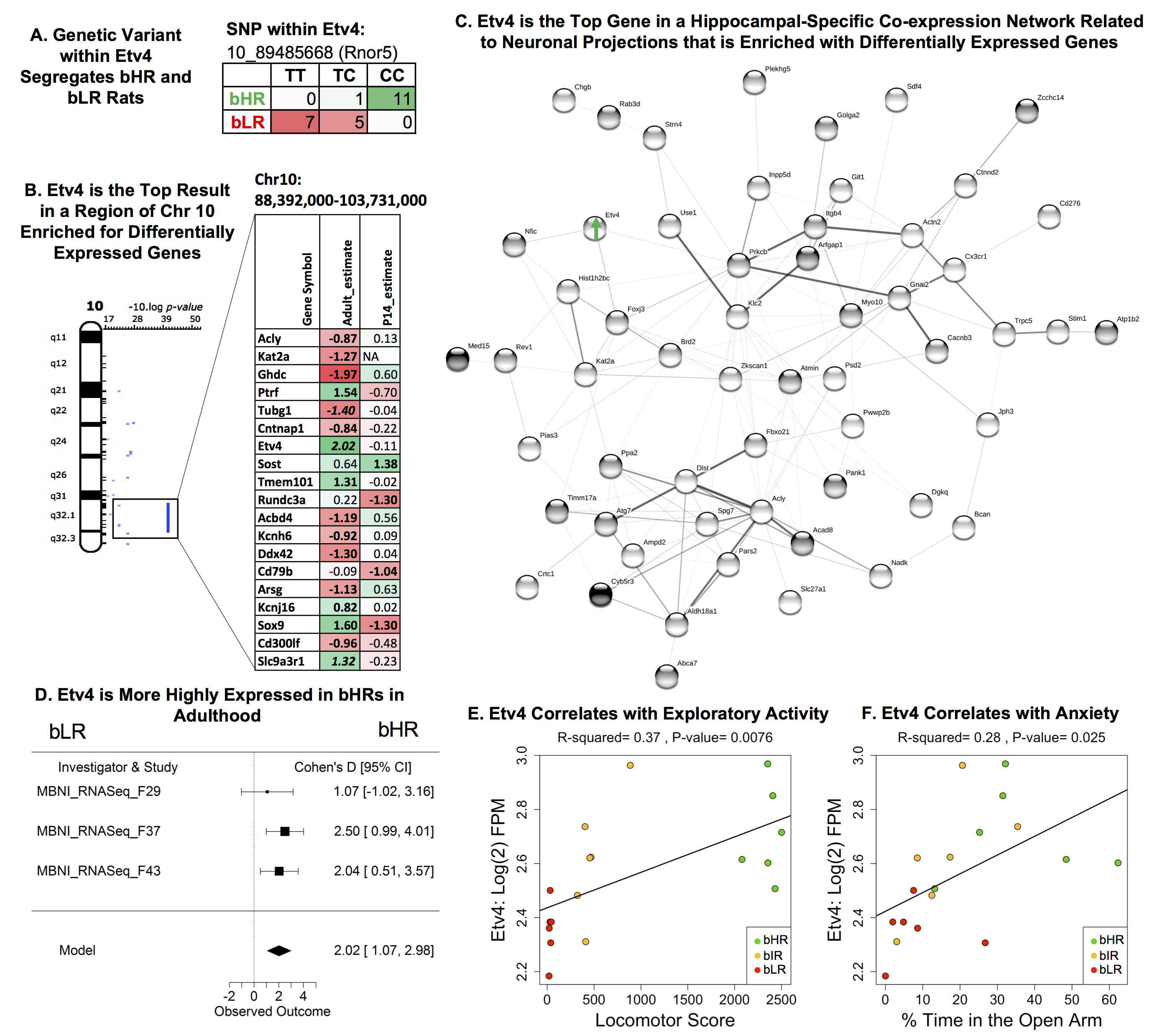

**Fig S8*.* ETS Variant 4 (Etv4) was the top gene within a hippocampal specific gene set and within a genomic loci on Chromosome 10 enriched for bHR/bLR differential expression. A)** Etv4 was the strongest result within a segment of Chromosome 10 enriched for differentially expressed genes that overlapped QTLs related to internalizing and externalizing behaviors (16, 25, 26). The table illustrates the differentially expressed genes within this region: estimate=estimated effect size (green/positive=more highly expressed in bHRs), bold=p<0.05, bold+italic=FDR<0.05. **B)** Etv4 was one of the top genes in a hippocampal specific co-expression network (*lightcyan*) that was enriched for bLR-upregulated genes in both development (FDR<0.05) and adulthood (FDR<0.10). Illustrated above is a PPI network (STRINGdb: confidence setting=0.40) constructed using the genes from this co-expression module that showed bHR/bLR differential expression in adulthood (n=74, p<0.05). This network was enriched with genes related to cell projections, neurons, synapses, and cation binding (FDR<0.05). **C)** A forest plot showing that Etv4 showed consistently higher expression in bHR rats (3 datasets, boxes=Cohen’s D from each study +/-95% confidence interval, “Model”=estimated effect size +/-95% confidence intervals provided by the meta-analysis, β=2.02, p=3.30E-05, FDR= 1.92E-02). **D-E)** In the behavioral data accompanying the MBNI_RNASeq_F37 dataset, Etv4 (units: log(2) fragments per million (FPM)) showed a positive relationship with **D)** exploratory locomotor activity (β=0.000131, R^2^= 0.37, p=0.0076) **E)** percent of time spent in the open arms of the elevated plus maze (β=0.00694, R^2^= 0.28, p=0.0249).

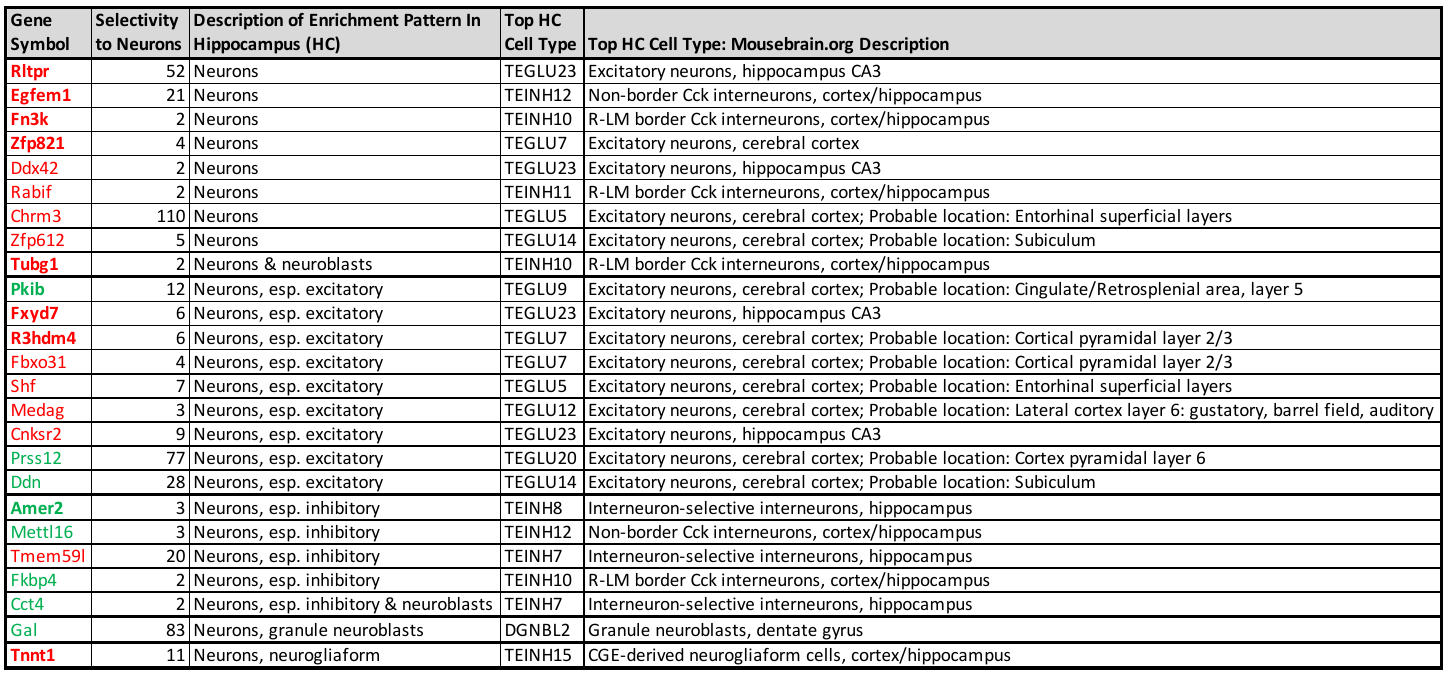

**Fig S9. Many of the top differentially expressed genes are predominantly expressed in particular hippocampal cell types, including neurons.** A comparison of our top meta-analysis results (FDR<0.10) with the new mousebrain.org database (Ziesel et al. 2018) suggested that many of our top genes (31%: 59/191 included in the analysis) have enriched (>2X) expression in a hippocampal cell type as compared to other hippocampal cell types. Shown above are the top bHR/bLR DE genes that have enriched expression in hippocampal neuronal cell types, which could not be properly interrogated within our cell type deconvolution analysis due to the dependence of *BrainInABlender* on cortically-derived datasets. The gene symbols are formatted to illustrate the results from our bHR/bLR meta-analysis (green=more highly expressed in bHRs, red=more highly expressed in bLRs, bold=FDR<0.05). “Selectivity to neurons” provides the ratio of the average expression value within the top hippocampal neuronal cell type vs. the top non-neuronal cell type. The hippocampal cell type with the most enriched expression is identified using the cluster name and description provided by mousebrain.org.
